# C9orf72/SMCR8 complex maintains microglial homeostasis via RAB8A-ESCRT-mediated lysosomal repair

**DOI:** 10.1101/2025.08.22.671707

**Authors:** Shan Li, Shidong Xu, Feng Li, Qirui Zhao, Penghui Zhang, Qinghua Guan, Xiangxiang Sun, Jundong Bi, Hu Xiao, Yiyuli Tang, Chen Peng, Qingfeng Chen, Yonghua Wang, Mei Yang

## Abstract

Microglia are critical regulators of neuroinflammation and neurodegeneration. Haploinsufficiency of C9orf72, the most common genetic cause of ALS and FTD, has been mainly linked to autophagy-lysosomal pathway defects, but its precise role in microglial lysosomal function remains unclear. Here we identify the C9orf72/SMCR8 complex as a key regulator of microglial homeostasis through lysosomal repair. Loss of C9orf72 and SMCR8 in mice causes age-dependent neuroinflammation and microgliosis, with microglia adopting a disease-associated state. In aged CNS tissue, microglia display lysosomal damage, marked by galectin-3 upregulation and its accumulation on lysosomes. To model this process, we applied the lysosomotropic agent LLOMe to microglia, which recapitulated lysosomal damage and revealed defective recruitment of phosphorylated RAB8A and the ESCRT machinery in C9orf72/SMCR8-deficient cells. Notably, mutant microglia accumulate GTP-bound RAB8A, which becomes aberrantly hyperphosphorylated and mislocalized to non-lysosomal vesicles. We further show that the GAP activity of the C9orf72/SMCR8 complex is essential for lysosomal repair. These findings uncover a previously unrecognized role for the C9orf72/SMCR8 complex in coordinating RAB8A-ESCRT-mediated lysosomal repair, thereby safeguarding microglial homeostasis and limiting neuroinflammation.

## Introduction

Microglial dysfunction represents a central pathogenic mechanism in neurodegenerative diseases, with aberrant activation states contributing to neuroinflammation and neuronal loss (Salter & Stevens, 2017). In neurodegenerative diseases, microglia adopt distinct activation states, including disease-associated microglia (DAM) and the microglial neurodegenerative phenotype (MGnD), characterized by altered gene expression profiles and enhanced phagocytic activity (Castro-Gomez & Heneka, 2024; Chen *et al*, 2024; Fumagalli *et al*, 2025; Keren-Shaul *et al*, 2017; Paolicelli *et al*, 2022). These activated microglia can contribute to neurodegeneration through excessive synaptic pruning, production of inflammatory mediators, and impaired clearance of protein aggregates (Heneka *et al*, 2025; Shi & Yong, 2025). Importantly, aging microglia exhibit compromised lysosomal function, characterized by accumulation of undegraded materials, elevated lysosomal pH, and increased susceptibility to membrane permeabilization. These age-related changes render microglia more prone to inflammatory activation and impair their neuroprotective functions(Quick *et al*, 2023).

Hexanucleotide repeat expansions in C9orf72 are the most common genetic cause of amyotrophic lateral sclerosis (ALS) and frontotemporal dementia (FTD) (DeJesus-Hernandez *et al*, 2011; Dobson-Stone *et al*, 2012; Rademakers, 2012; Sha *et al*, 2012). In addition to toxic gain-of-function effects, C9orf72 haploinsufficiency has emerged as a critical driver of disease progression (Boivin et al, 2020; Shao et al, 2019; Zhu et al, 2020). C9orf72 forms a heterotrimeric complex with SMCR8(Smith-Magenis syndrome chromosome region candidate 8) and WDR41(WD repeat-containing protein 41), which regulates diverse cellular processes including autophagy, endo-lysosomal pathways, and membrane trafficking. (Amick *et al*, 2016; Sellier *et al*, 2016; Shi *et al*, 2018; Su *et al*, 2020; Sullivan *et al*, 2016; Tang *et al*, 2020; Ugolino *et al*, 2016; Yang *et al*, 2016; Zhang *et al*, 2023; Zhang *et al*, 2018). Structural analyses have shown that this complex exhibits GTPase-activating protein (GAP) activity toward multiple RAB GTPases, including RAB8A(Su *et al*., 2020; Tang *et al*., 2020; Tang *et al*, 2023).

Evidence from human iPSC-derived models, mouse genetics, and patient tissue consistently demonstrates that C9orf72, together with SMCR8, is essential for immune homeostasis. At the systemic level, C9orf72 or SMCR8 deficiency results in splenomegaly, lymphadenopathy, autoimmunity, and premature mortality, phenotypes linked to dysregulated mTORC1–AKT signaling, excessive lysosomal exocytosis, and prolonged TLR signaling (Lan *et al*, 2019; McAlpine *et al*, 2018; O’Rourke *et al*, 2016; Shao *et al*, 2020; Zhang *et al*., 2018). In myeloid cells (macrophages, monocytes, microglia, dendritic cells), loss of C9orf72 disrupts lysosomal function, impairs autophagy initiation, and triggers aberrant activation of STING-type I interferon and JAK/STAT signaling, while also enhancing antigen presentation and CD80 upregulation. (Burberry *et al*, 2020; Lall *et al*, 2021; Limone *et al*, 2024; McCauley *et al*, 2020; O’Rourke *et al*., 2016; Pang & Hu, 2023; Vahsen *et al*, 2023). In microglia, C9orf72 deficiency drives a transition from a homeostatic to inflammatory state, leading to excessive release of MMP9, abnormal synaptic pruning and non–cell-autonomous neurotoxicity that contributes to cognitive deficits (Banerjee *et al*, 2023; Vahsen *et al*., 2023; Wang *et al*, 2022) .

Although inflammatory phenotypes of *C9orf72/SMCR8*-deficient microglia are well established, the upstream mechanisms by which this complex safeguards microglial homeostasis remain unclear. Lysosomal dysfunction has emerged as a convergent driver of age-related and neurodegenerative processes(Ferrari *et al*, 2024). In aging microglia, lysosomes accumulate undegradable substances including lipofuscin, oxidized proteins, and iron, leading to elevated lysosomal pH, membrane permeabilization, and inflammatory activation(Quick *et al*., 2023). Upon lysosomal membrane permeabilization, cells engage quality control pathways, including galectin sensing, ESCRT-mediated repair, lysophagy, and additional mechanisms such as ER-mediated lipid transfer, sphingomyelinase-driven ceramide generation, annexin recruitment, and stress granule stabilization (Radulovic *et al*, 2025). In macrophages, RAB8A phosphorylation by LRRK2 facilitates ESCRT recruitment (Eguchi *et al*, 2018; Herbst *et al*, 2020). Despite these advances, the precise molecular mechanisms by which C9orf72/SMCR8 deficiency leads to microglial dysfunction remain incompletely understood. Given the emerging role of lysosomal membrane damage and repair in microglial aging, we hypothesized that C9orf72/SMCR8 complex regulates microglial homeostasis through control of lysosomal repair pathways.

Here, using knockout mouse models and cellular systems, we demonstrate that C9orf72/SMCR8 deficiency results in age-dependent microglial activation and neuroinflammation. Mechanistically, we show for the first time that loss of this complex disrupts ESCRT recruitment and alters RAB8A phosphorylation dynamics, thereby impairing lysosomal repair. These findings establish a previously unrecognized role of the C9orf72–SMCR8 complex in microglial lysosomal quality control and provide mechanistic insight into how its loss contributes to neuroinflammation in ALS/FTD. **Results**

### Loss of *C9orf72* and *Smcr8* leads to age-dependent neuroinflammation and microgliosis in mice

To investigate the role of C9orf72 and SMCR8 in microglia, we first examined the expression levels of these proteins in various cell types of the central nervous system (CNS). Analysis of single cell RNA-seq data from the *Whole Cortex & Hippocampus - 10x genomics*(Yao *et al*, 2021) dataset revealed that C9orf72 and SMCR8 are highly expressed in microglia compared to other CNS cell types (Figure S1A). Western blot analysis demonstrated differential expression of C9orf72 and SMCR8, with higher protein levels detected in primary microglia compared to neurons isolated from wild-type mice (Figure S1B, C). Western blot analysis across age groups showed a modest increase in C9orf72 and SMCR8 protein levels in 20-month-old mice compared to 4- and 12-month-old animals in the brain and spinal cord (Figure S1D, E, F). This age-dependent upregulation suggests that the C9orf72-SMCR8 complex may play an increasingly important role in maintaining microglial homeostasis during aging.

To elucidate the functional consequences of *C9orf72* and *Smcr8* loss, we generated double knockout (dKO) mice (Figure S1D). Immunohistochemical analysis revealed significantly increased IBA1-positive activated microglia in the hippocampus and spinal cord of dKO mice compared to age-matched WT controls (Figure 1A). This microgliosis phenotype intensified with age, suggesting an age-dependent effect of C9orf72 and SMCR8 deficiency. To further investigate the inflammatory status of dKO mice, we measured the mRNA levels of pro-inflammatory cytokines in the cortex and spinal cord of 4-month-old and 20-month-old mice. We observed significant upregulation of *Tnf* and *Il1b* in the brain, and *Il1b* and *Il6* in the spinal cord of dKO mice compared to heterozygous controls, with the effect being more pronounced in older mice (Figure 1B, C). Immunofluorescence staining for neuronal markers revealed significantly decreased numbers of NeuN- and ChAT-positive neurons in both hippocampus and spinal cord of 20-month-old dKO mice compared to age-matched WT mice, indicating age-dependent neurodegeneration (Figure 1D-G). These results demonstrate that loss of *C9orf72* and *Smcr8* leads to age-dependent neuroinflammation, microgliosis and neurodegeneration.

**Figure 1.**
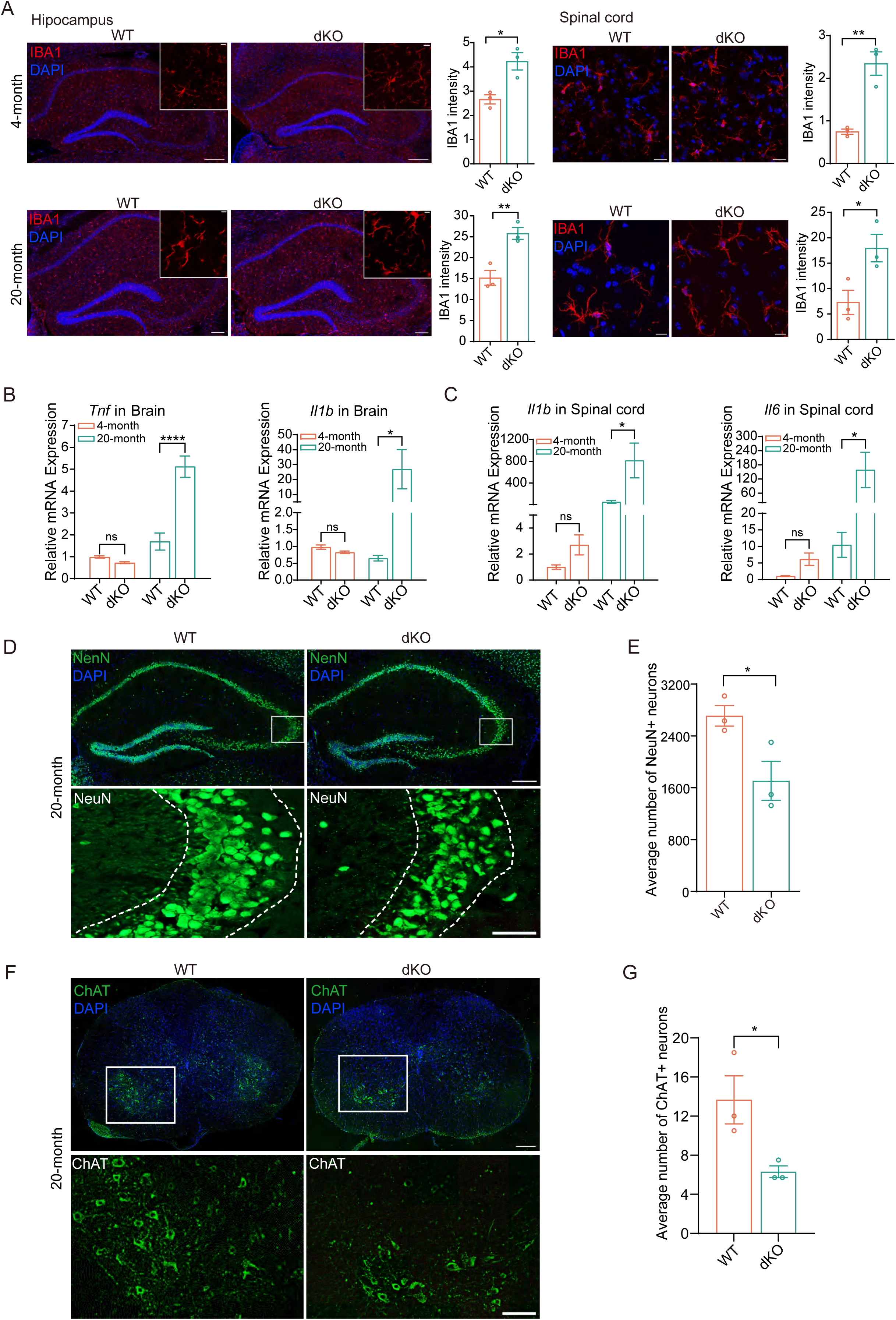
*C9orf72*/*Smcr8* double knockout mice develop age-dependent neuroinflammation and neurodegeneration. (A) Representative Immunofluorescence images of IBA1 (red) and DAPI (blue) in hippocampus (scale bar, 200 µm) and spinal cord (scale bar, 20 µm) from WT and dKO mice at 4 and 20 months of age. Insets, higher magnification (scale bar, 10 µm). Quantification of IBA1 intensity shown on right. (B) qRT-PCR analysis of *Tnf* and *Il1b* mRNA expression in brain tissue at 4 and 20 months of age. Data normalized to 4-month WT controls. (C) qRT-PCR of *Il1b* and *Il6* mRNA expression in spinal cord tissue at 4 and 20 months. (D, E) Representative NeuN immunostaining (green) and DAPI (blue) in hippocampus of 20-month-old WT and dKO mice. Scale bar: 200 μm(overview), 50 μm (insets). Quantification of NeuN positive neurons in hippocampus shown in (E). (F, G) Representative ChAT immunostaining (green) and DAPI (blue) in spinal cord sections from 20-month-old WT and dKO mice. Scale bars: 200 μm (overview), 100 μm (insets). Quantification of ChAT-positive motor neurons in spinal cord shown in (G). Data are mean ± SEM (n = 3 mice per group from three independent experiments). Student’s unpaired two-tailed t-test (A, E, G) or two-way ANOVA with Bonferroni post-test (B, C). *P < 0.05, **P < 0.01, ****P < 0.0001; ns, not significant.

Recent studies have identified distinct microglial phenotypes in neurodegenerative conditions, including disease-associated microglia (DAM) and the microglial neurodegenerative phenotype (MGnD), which share overlapping characteristics. Both phenotypes share common molecular and functional characteristics, suggesting they may represent overlapping states of microglial activation in response to pathological conditions. To gain deeper insights into the microglial response, we performed single-nucleus RNA sequencing (snRNA-seq) on brains from aged (20- to 24-month-old) WT and dKO mice. After stringent quality control including doublet removal, we obtained 33,257 high-quality single-cell transcriptomes (WT: n=16,707 cells; dKO: n=16,550 cells from n=3 mice per genotype) that were annotated into major cell types based on cell-type-specific markers (Figure 2A). Cell-type composition analysis revealed a significant expansion of the microglial population in dKO mice compared to WT controls (Figure 2B). UMAP visualization demonstrated distinct clustering of major CNS cell types including neurons, astrocytes, oligodendrocytes, and microglia (Figure 2A, C). Further analysis identified differentially expressed genes (DEGs) in dKO microglia compared to WT, with significant upregulation of DAM/MGnD markers including *Apoe*, *Gpnmb*, *Lgals3(gal3)*, *Trem2* and *Clec7a* (Figure 2D). Gene Ontology (GO) analysis of upregulated genes revealed enrichment of pathways involved in innate immune response, reactive oxygen species metabolic processes, regulation of lymphocyte proliferation, canonical inflammasome complex assembly, and cytokine production (Figure 2E). Consistent with the snRNA-seq data, qPCR analysis of isolated primary microglia confirmed significant upregulation of these DAM-associated genes in dKO mice (Figure 2F). This molecular signature suggests that loss of *C9orf72* and *Smcr8* leads to acquisition of a disease-associated microglial phenotype characterized by enhanced inflammatory responses and altered cellular homeostasis.

**Figure 2.**
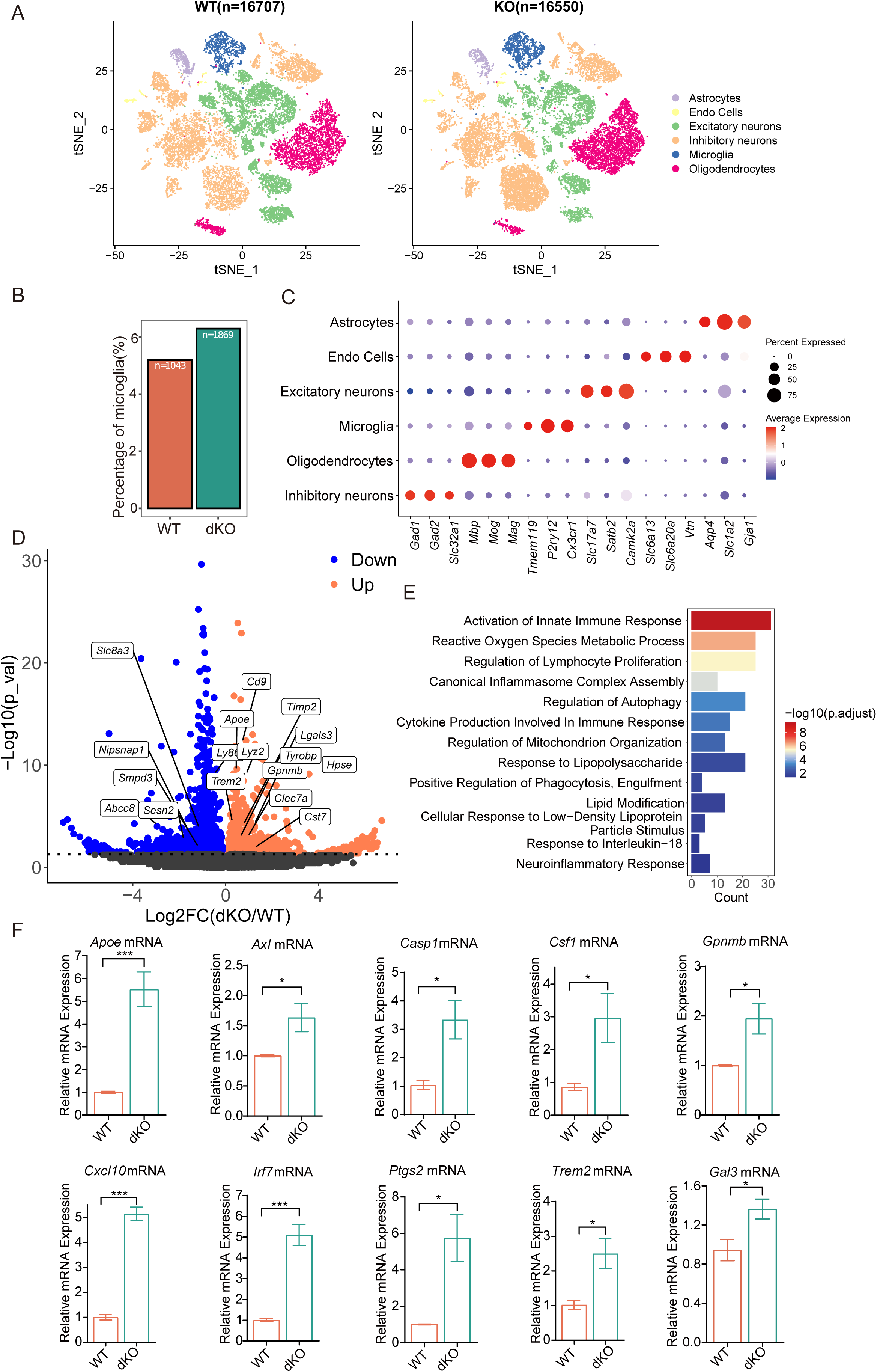
snRNA-seq and primary microglia qRT-PCR reveal disease-associated microglia (DAM) activation in dKO mice. (A) UMAP of 33,257 single-nucleus transcriptomes from brains of 20- to 24-month-old WT and dKO mice (WT: n = 16,707 cells; dKO: n = 16,550 cells from 3 mice per genotype) mice. Major CNS cell types annotated by marker gene expression. (B) Quantification of microglial cell proportion among total cells in WT and dKO samples. (C) Dot plot showing expression of selected cell-type-specific marker genes across identified cell populations. Circle size indicates percentage of expressing cells; color intensity represents average normalized expression level. (D) Volcano plot showing differentially expressed genes (DEGs) in dKO versus WT microglia. Upregulated DAM-associated markers are highlighted. (E) Gene Ontology (GO) enrichment of biological processes for upregulated genes in dKO microglia. Color scale represents -log10 adjusted P-value. (F) qRT-PCR validation of DAM-associated genes (*Apoe, Axl, Casp1, Csf1, Gpnmb, Cxcl10, Irf7, Ptgs2, Trem2, Gal3)* in primary microglia from WT and Dko mice. Data normalized to Actin expression. Data are mean ± SEM (n = 3 mice per group from three independent experiments). Student’s unpaired t-test for qRT-PCR. *P < 0.05, ***P < 0.001.

### Age-dependent inflammation correlates with increased levels of Galectin-3

Our snRNA-seq analysis revealed significant upregulation of Galectin-3 (*Gal3*) mRNA expression in microglia from dKO mice, prompting us to investigate the role of GAL3 in the observed neuroinflammatory phenotype. Given that GAL3 serves as a key marker for lysosomal dysfunction and microglial inflammatory responses(Jia *et al*, 2020; Shan *et al*, 2024; Siew *et al*, 2019), several research indicates lysosome repair lined to neurodegeneration(Chou *et al*, 2025; Radulovic *et al*., 2025). We investigated whether galectin family members were altered in dKO mice. Western blot analysis of brain tissues revealed a progressive age-dependent increase in GAL3 protein levels in dKO mice compared to WT controls, with significant elevation observed in both cortex and spinal cord (Figure 3A). In contrast, GAL8 and GAL9 showed no significant differences between dKO and WT mice (Figure S2D, E). Consistent with the protein data, quantitative RT-PCR analysis confirmed significant elevation of *Gal3* mRNA levels in both brain and spinal cord of 20-month-old dKO mice (Figure 3B, C). To visualize GAL3’s subcellular distribution *in vivo*, we performed immunofluorescence staining using GAL3 antibodies on cortical and spinal cord sections from 20-month-old WT and dKO mice. We observed a higher proportion of GAL3-positive, CD68-positive microglia in the mutant (dKO) mice, with clear colocalization (Figure 3D-G). Additionally, we administered AAV-PHP.eB-GAL3-GFP reporter vectors via tail vein injection to achieve broad CNS expression. Remarkably, dKO mice exhibited increased GAL3-GFP puncta in IBA1-positive microglia, with colocalization with the lysosomal marker LAMP1 (Figure 3H, J). Quantification revealed significantly higher percentage of microglia containing GAL3^+^LAMP1^+^ lysosomes in dKO mice compared to WT, in both brain and spinal cord (Figure 3H-K). These results suggest that the age-dependent inflammation observed in dKO mice correlates with increased levels of GAL3, potentially indicating an accumulation of damaged lysosomes in mutant microglia.

**Figure 3.**
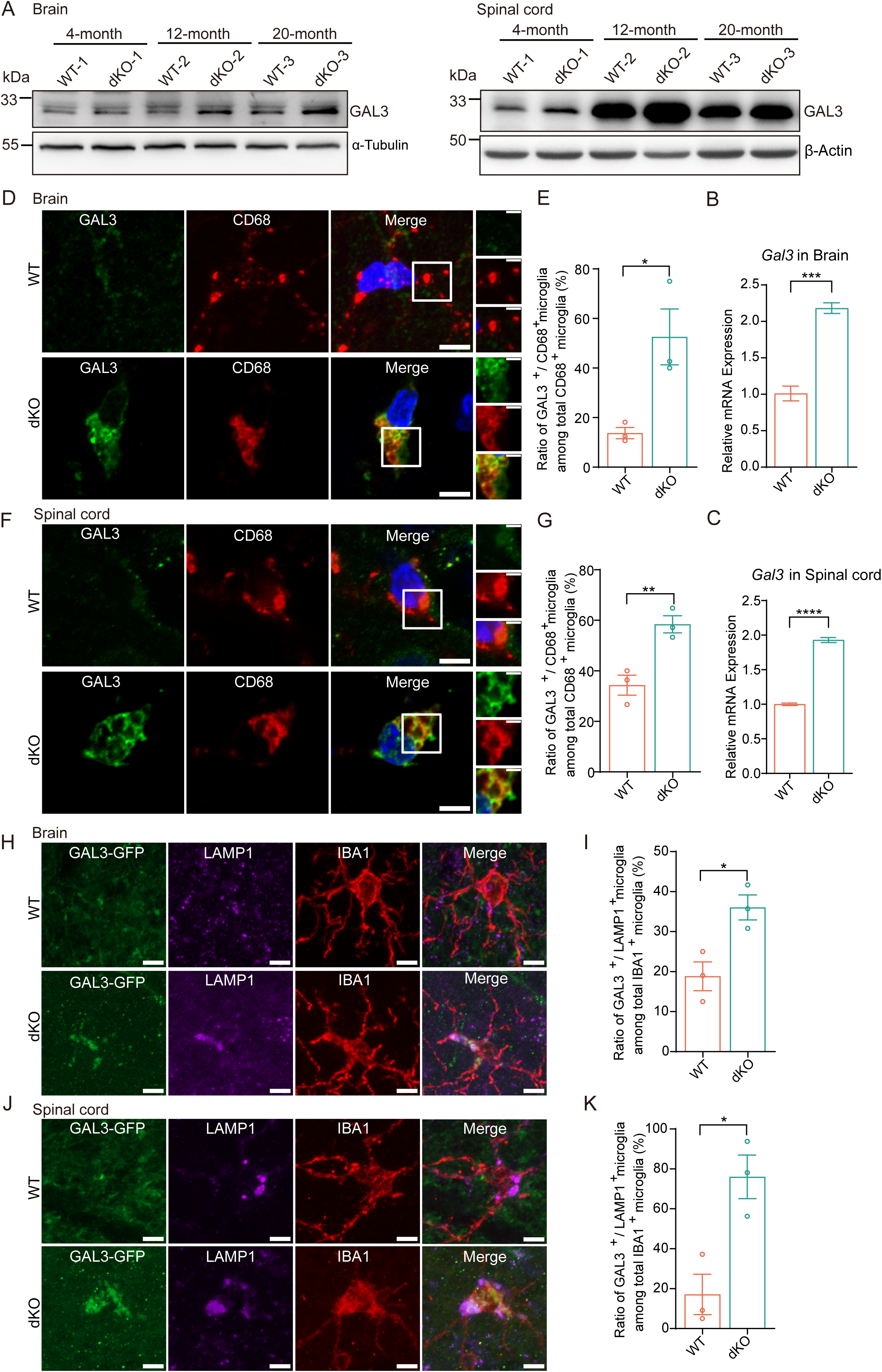
Age-dependent GAL3 upregulation and enhanced lysosomal localization in *C9orf72/Smcr8* dKO mice. (A) Western blot analysis of GAL3 protein levels in brain and spinal cord tissue at 4, 12, and 20 months of age. α-Tubulin and β-actin serve as loading controls. (B, C) qRT-PCR analysis of *Gal3* mRNA expression in brain (B) and spinal cord (C) from 20-month-old WT and dKO mice. (D, F) Representative immunofluorescence images of GAL3 (green), CD68 (red), and DAPI (blue) in brain cortex (D) and spinal cord (F) sections from 20-month-old mice. Scale bars: 5 µm (overview), 2 µm (insets). (E, G) Quantification of GAL3-positive/CD68-positive microglia in cortex (E) and spinal cord (G). Each data point represents the mean of 3-5 fields from 3 sections per mouse. (H, J) Representative images showing GAL3-GFP (green), LAMP1 (purple), and IBA1 (red) expression in cortex (H) and spinal cord (J) following AAV-PHP.eB delivery. Scale bar: 5 μm. (I, K) Quantification of the percentage of microglia (Iba1) harboring GAL3-GFP-positive lysosomes (LAMP1) in the cortex (I) and spinal cord (K). Data presented as mean ± SEM (n = 3 mice per group from three independent experiments). Statistical analysis by Student’s two-tailed unpaired t-test. *P < 0.05, ***P < 0.001 and ****P <0.0001.

### C9orf72/SMCR8 deficiency enhances stress-induced inflammatory and phagocytic responses in microglia

To investigate how C9orf72/SMCR8 deficiency affects microglial function, we isolated primary microglia from WT and KO mice and examined their inflammatory responses. Under basal conditions, primary microglia from dKO mice showed comparable levels of *Tnf*, *Il6*, and *Il1β* to WT mice. Upon lipopolysaccharide (LPS) and interferon-γ (IFN-γ) stimulation, which promotes classical microglial activation, dKO primary microglia exhibited significantly elevated expression of these pro-inflammatory cytokines (Figure 4A), consistent with previous findings that C9orf72 deficiency leads to enhanced pro-inflammatory responses in myeloid cells (Lall *et al*., 2021; Limone *et al*., 2024; Pang & Hu, 2023). To further investigate lysosomal damage-induced inflammation in a cell line model, we generated *C9orf72* and *Smcr8* single and double knockout BV2 cell lines using CRISPR-Cas9 technology (Figure S2A-C). When challenged with lysosomal stress induced by 1mM LLOMe treatment for 30 minutes, *C9orf72* KO BV2 cells showed comparable *Il1b* expression to WT cells. However, after LLOMe treatment followed by a 6-hour recovery period (washout), both single knockouts (*C9orf72* KO and *Smcr8* KO) and dKO cells exhibited significantly enhanced *Il1b* expression compared to WT cells (Figure 4B). Notably, primary microglia demonstrated markedly greater sensitivity to lysosomal damage-induced inflammatory responses compared to the BV2 cell line model. We validated these findings using primary microglia, where dKO cells displayed significantly increased *Tnf* and *Il1b* expression specifically after 30-minute LLOMe treatment compared to WT cells (Figure 4B), suggesting enhanced inflammatory responses upon lysosomal damage. This heightened inflammatory response suggests a critical role for C9orf72 and SMCR8 in maintaining lysosomal integrity and regulating inflammatory signaling pathways upon lysosomal damage.

**Figure 4.**
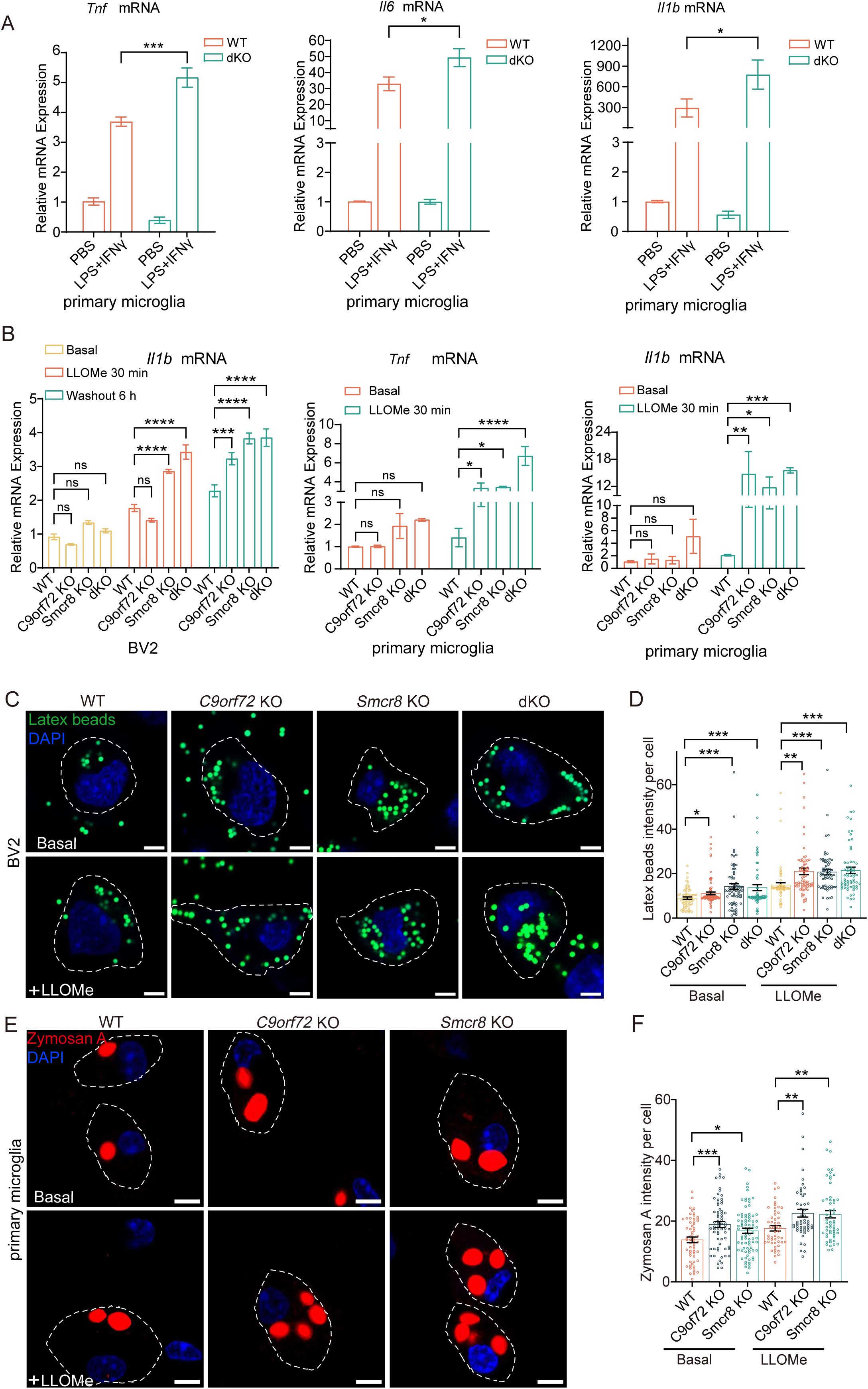
*C9orf72/Smcr8* deficiency enhances stress-induced microglial hyperactivation and enhanced phagocytosis. (A) qRT-PCR analysis of *Tnf*, *Il6*, and *Il1b* mRNA expression in primary microglia from WT and dKO under basal conditions or LPS (100 ng/ml)/IFN-γ(50 ng/ml) stimulation for 24 hours. (B) Cytokine mRNA expression following LLOMe treatment. BV2 cells: basal, 30 min LLOMe (1 mM), and 6 h washout conditions. Primary microglia: basal and 30 min LLOMe (0.5 mM) treatment. (C, D) Latex bead phagocytosis in BV2 cells. (C) Representative images showing internalized beads (green) and nuclei (DAPI, blue). Scale bar, 5 µm. (D) Quantification of internalized beads intensity. (E, F) Zymosan A bioparticle uptake in primary microglia. (E) Representative images showing internalized zymosan particles (red) and nuclei (DAPI, blue). Scale bar: 5 μm. (F) Quantification of particle intensity per cell. Data presented as mean ± SEM (n = 3 biological replicates per group from three independent experiments). For phagocytosis assays, ≥60 cells were analyzed per genotype per condition across three independent experiments (≥20 cells per experiment). Statistical analysis by two-way ANOVA followed by Bonferroni post-test (A, B) or Student’s two-tailed unpaired t-test (D, F): *P < 0.05, **P < 0.01, *** < 0.001, ****P < 0.0001, ns: not significant.

Phagocytosis is a key function of microglia, allowing them to clear apoptotic cells, protein aggregates, and other debris from the CNS. Using latex beads uptake assays in BV2 cells, we found that both single and double knockout cells exhibited increased phagocytic activity under basal conditions. This enhanced phagocytic capacity was further enhanced following 1mM LLOMe treatment (Figure 4C, D). Similarly, when assessing primary microglia using zymosan A particles, both *C9orf72* KO and *Smcr8* KO cells showed enhanced phagocytic activity at baseline, with further increases observed after 0.5 mM LLOMe treatment (Figure 4E, F). These results demonstrate that loss of either *C9orf72* or *Smcr8* leads to dysregulated microglial responses, particularly under stress conditions, suggesting their cooperative role in maintaining proper microglial function.

### *C9orf72/Smcr8* loss of function exacerbates lysosomal damage and impairs recovery following LLOMe treatment

Given our observations of enhanced inflammatory and phagocytic responses in C9orf72/SMCR8-deficient microglia, we investigated the subcellular localization and function of C9orf72/SMCR8 during lysosomal stress. Immunofluorescence analysis revealed that C9orf72 significantly colocalized with LAMP1 upon lysosomal damage induced by LLOMe treatment (Figure S3A). Using the TMEM192-3xHA lysosomal immunoprecipitation system, we further confirmed enhanced association of both C9orf72 and SMCR8 with lysosomes during lysosomal stress conditions (Figure S3B), suggesting their potential roles in lysosomal stress response. To directly assess lysosomal function, we performed LysoTracker staining in BV2 cells. Under basal conditions, all genotypes showed comparable LysoTracker signals. Following 1mM LLOMe treatment, WT cells showed initial decrease in LysoTracker signal followed by recovery after 3-hour washout. In contrast, both single knockout and dKO cells exhibited markedly slower recovery, with LysoTracker signal remaining consistently and significantly lower than that of WT cells throughout the observation period, indicating impaired lysosomal repair and recovery following damage (Figure 5A, B). To further characterize lysosomal damage, we examined GAL3 that specifically binds to β-galactoside residues exposed upon lysosomal membrane permeabilization and damage. Immunofluorescence analysis revealed significantly increased GAL3-positive lysosomes in both single and double knockout cells. This increase was evident during 30-60 minutes of LLOMe treatment. The enhanced GAL3 puncta persisted even after LLOMe washout (at 3-, and 6-hours post-washout), indicating prolonged lysosomal damage in mutant cells (Figure 5C, D). Lyso-IP analysis confirmed increased lysosome-associated GAL3 in mutant cells (Figure 5E). Consistently, primary microglia from single and double knockout mice showed markedly elevated GAL3 levels as early as 30 minutes post-LLOMe (0.5mM) treatment (Figure S3C D). In BV2 cells, *C9orf72* KO, *Smcr8* KO, and dKO cells showed significantly reduced mature CTSD levels, indicating defective lysosomal protease. (Figure S3E). Electron microscopy analysis provided further visual evidence of lysosomal dysfunction in mutants. We categorized lysosomes as “defective lysosomes” based on the presence of characteristic morphological alterations including lysosomal enlargement, decreased electron density within the lysosomal lumen, and visible membrane ruptures. We observed a significant increase in the number of visibly defective lysosomes in LLOMe-treated mutant cells (Figure 5 F, G and Figure S3F, G). These findings suggest that C9orf72 and SMCR8 play critical roles in maintaining lysosomal homeostasis, and their loss leads to compromised lysosomal function and impaired recovery from lysosomal damage.

**Figure 5.**
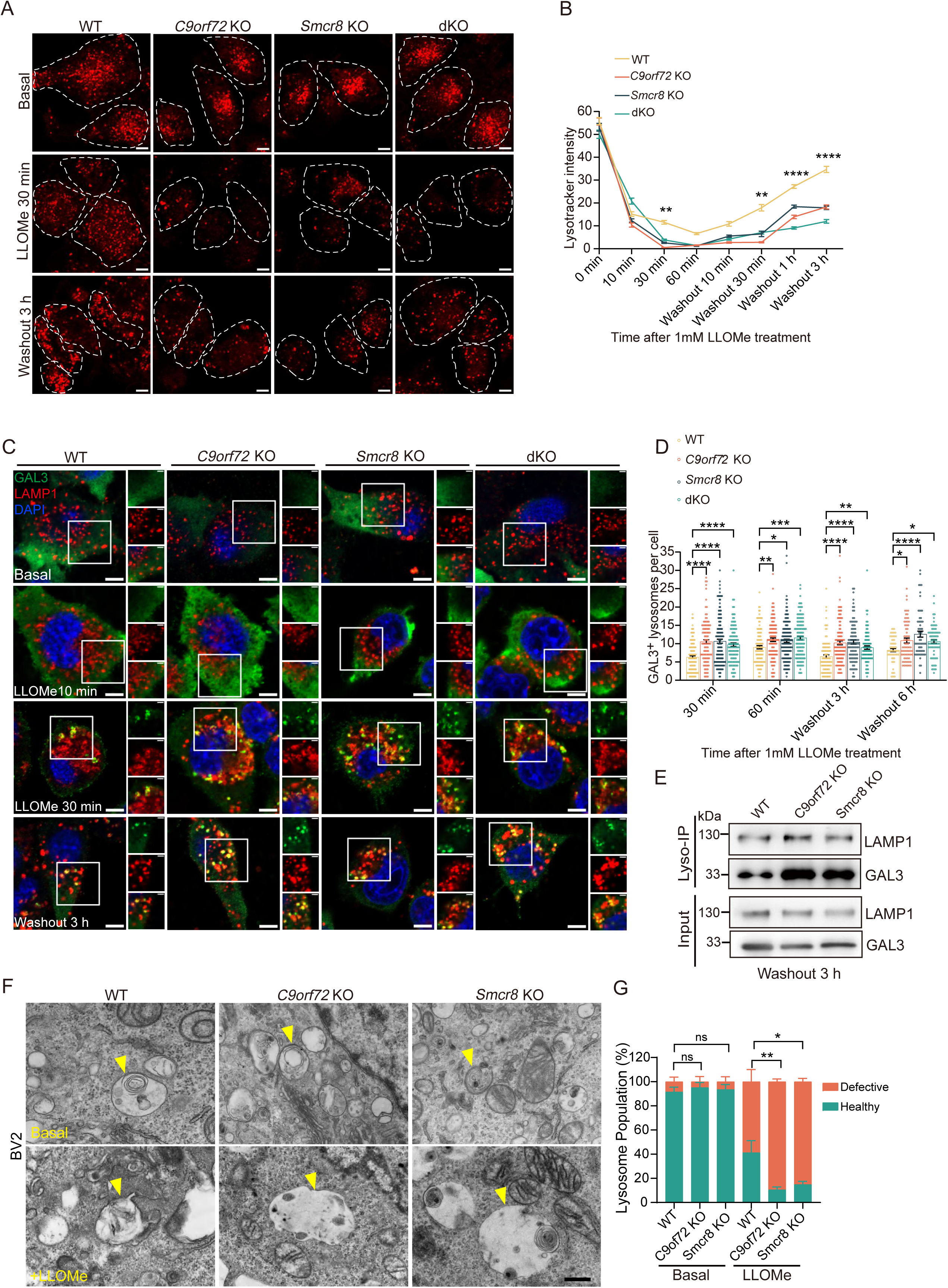
*C9orf72/Smcr8* deficiency impairs lysosomal membrane repair and increases lysosomal damage. (A) Representative fluorescence images of LysoTracker Red in WT, C9orf72 KO, Smcr8 KO, and dKO BV2 cells under basal conditions, after 30 min LLOMe (1 mM) treatment, and following 3 h washout. Nuclei stained with DAPI (blue). Dashed lines indicate cell boundaries. Scale bar: 5 μm. (B) Time-course quantification of LysoTracker Red fluorescence intensity following LLOMe treatment and washout. (C) Immunofluorescence of GAL3 (green), LAMP1 (red), and DAPI (blue) at indicated times after LLOMe. Scale bar: 5 µm. (D) Quantification of GAL3-positive lysosomes per cell across time points. (E) LysoIP using TMEM-192-3xHA system from WT, *C9orf72* KO, and *Smcr8* KO BV2 cells at 3 h post-LLOMe washout. Input and IP fractions analyzed by Western blot for GAL3 and LAMP1. (F) TEM images of BV2 cells under basal and 30 min post-LLOMe treatment. Arrowheads indicate lysosomes. Scale bar: 500 nm. (G) Quantification of defective lysosomes (% of total). Defects defined by lysosomal enlargement, reduced electron density, or membrane rupture. Data are shown as mean ± SEM from three independent experiments. For cellular analyses, ≥60 cells were analyzed per genotype per condition across three independent experiments (≥20 cells per experiment). Statistical analysis by two-way ANOVA followed by Bonferroni post-test (B, D) and unpaired two-tailed t-test (G). *P < 0.05, **P < 0.01, ***< 0.001****P < 0.0001, ns: not significant.

### Impaired recruitment of ESCRT machinery to damaged Lysosomes in C9orf72/SMCR8-deficient cells

Since ESCRT machinery is rapidly recruited to repair damaged lysosomes, acting as a primary defense mechanism that precedes autophagy-mediated clearance (lysophagy)(Radulovic *et al*, 2018; Skowyra *et al*, 2018), we examined whether ESCRT recruitment was compromised in *C9orf72/Smcr8*-deficient cells. Lyso-IP experiments revealed significantly reduced association of ALIX (ALG-2-interacting protein X) with lysosomes in mutant cells (Figure 6A). ALIX functions as an ESCRT-associated adaptor protein that bridges ESCRT-I and ESCRT-III complexes to facilitate membrane scission during lysosomal repair processes(Radulovic *et al*., 2018). To further characterize ESCRT recruitment dynamics, we examined CHMP2B recruitment to lysosomes using immunofluorescence analysis. Under basal conditions, CHMP2B showed diffuse cytoplasmic distribution across all genotypes. Following 1mM LLOMe treatment, WT cells exhibited increased CHMP2B puncta that recruited to lysosomes (LAMP1 positive vesicles) within 10-30 minutes. In contrast, both single and double knockout cells showed significantly reduced CHMP2B-positive lysosomes at these early time points (Figure 6B, C). These findings suggest that the C9orf72/SMCR8 complex plays a crucial role in facilitating ESCRT-mediated lysosomal repair.

**Figure 6.**
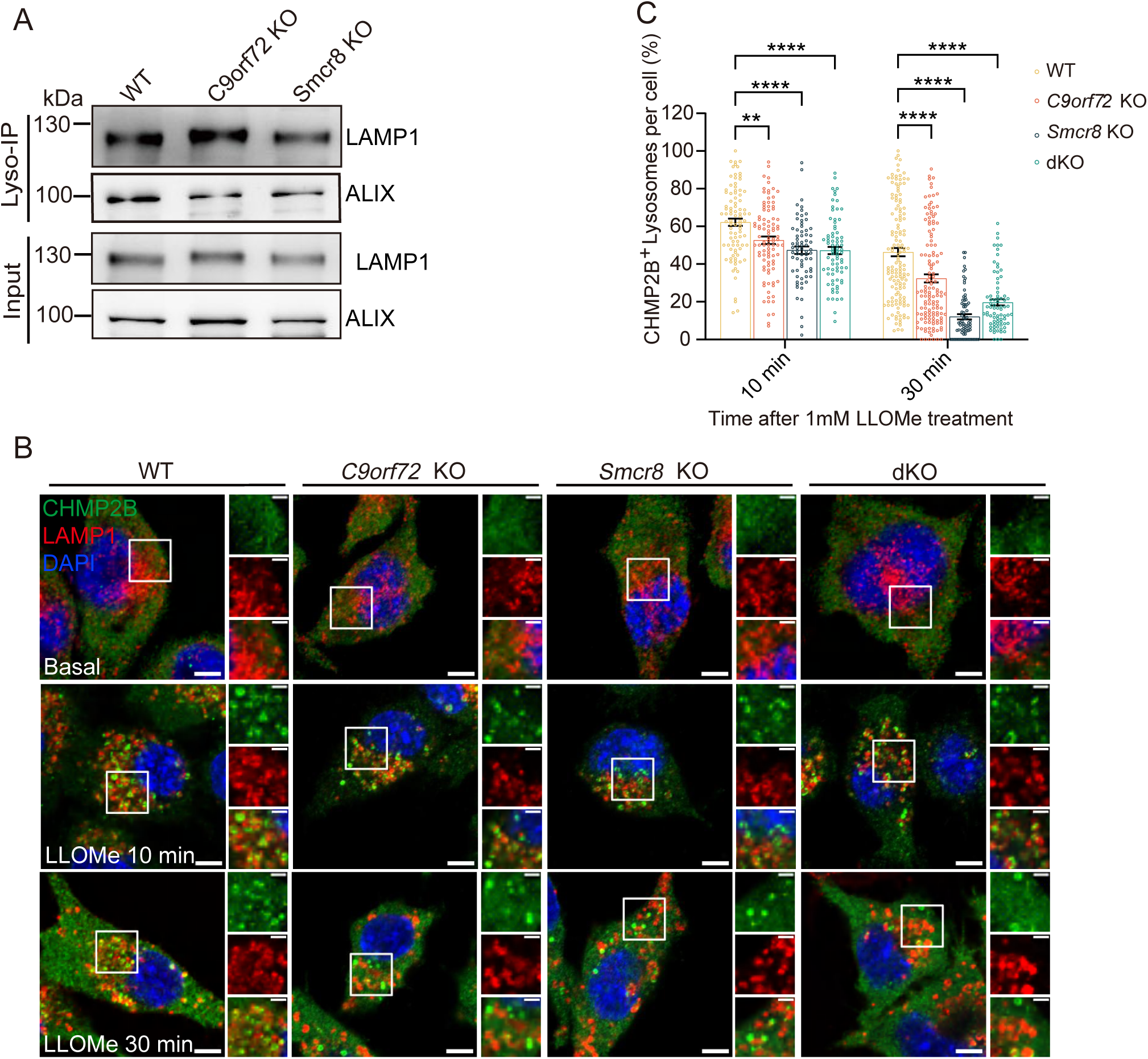
C9orf72 and SMCR8 deficiency impairs ESCRT recruitment. (A) LysoIP analysis of WT, *C9orf72* KO, and *Smcr8* KO BV2 cells. Total lysates and LysoIP fractions analyzed by Western blot for ALIX and LAMP1. (B) Representative immunofluorescence images of CHMP2B (green), LAMP1 (red), and DAPI (blue) in WT and knockout BV2 cells under basal conditions and after LLOMe treatment (1 mM, 10 and 30 min). Scale bars: 5 μm (overview), 2 μm (insets). (C) Quantification of CHMP2B recruitment to lysosomes, expressed as percentage of LAMP1-positive lysosomes showing CHMP2B colocalization per cell. Each dot represents an individual cell. Data are presented as mean ± SEM from three independent experiment. For immunofluorescence quantification, ≥70 cells were analyzed per genotype per condition across three independent experiments. Statistical analysis was performed using Two-way ANOVA followed by Bonferroni post-test. **P < 0.01, ****P < 0.0001.

### C9orf72/SMCR8 deficiency reduces lysosomal phospho-RAB8A while increasing cytoplasmic puncta

Phosphorylated RAB8A (pT72-RAB8A), activated by LRRK2 kinase, plays a critical role in recruiting ESCRT-III components to damaged lysosomes for membrane repair in macrophages(Herbst *et al*., 2020). To investigate how C9orf72/SMCR8 deficiency affects this process, we examined phospho-RAB8A dynamics following LLOMe-induced lysosomal damage in BV2 cells. In WT cells under basal conditions, phospho-RAB8A displayed minimal punctate staining with limited lysosomal association. Following 1mM LLOMe treatment, we observed rapid and dramatic recruitment of phospho-RAB8A to LAMP1-positive lysosomes within 30 minutes of exposure (Figure S4A). The dramatic shift from this diffuse, non-lysosomal distribution to prominent lysosomal recruitment upon LLOMe treatment demonstrates the specificity of the damage-induced response. Western blot analysis revealed a time-dependent increase in phospho-T72 RAB8A levels, initiating at 30 minutes of LLOMe treatment. Phospho-RAB8A signals persisted even after LLOMe washout, remaining detectable at 3 hours and 6 hours post-washout (Figure S4B). *C9orf72/Smcr8*-deficient cells exhibited markedly altered phospho-RAB8A distribution patterns. While wild-type cells showed efficient pT72-RAB8A recruitment to LAMP1-positive lysosomes, knockout cells displayed significantly reduced lysosomal localization of phospho-RAB8A. Interestingly, total phospho-RAB8A puncta increased in mutant cells, with enhanced accumulation in non-lysosomal cytoplasmic compartments (Figure 7A-D). Western blot analysis further revealed distinct recovery kinetics between genotypes. Both single knockout and dKO cells displayed significantly elevated pT72-RAB8A levels compared to WT cells across all time points examined, including 30-minute and 60-minute LLOMe treatment, as well as after a 3-hour washout period (Figure 7 E-G). The phosphorylation levels peaked at 60 minutes of LLOMe treatment in the knockout cells, suggesting that lysosomal damage repair defects in mutant cells are accompanied by hyperphosphorylation of RAB8A, along with a modest increase in total RAB8A protein levels. This phenotype was recapitulated *in vivo,* with elevated pT72-RAB8A signals detected in brain and spinal cord tissues from 20-month-old knockout mice (Figure 7H-M), indicating that the dysregulated RAB8A phosphorylation observed in cell culture models reflects pathological changes occurring in the aging nervous system. We observed that the increased phosphorylation signal was accompanied by slightly elevated total RAB8A protein levels, suggesting that lysosomal damage triggers both enhanced RAB8A phosphorylation and protein stabilization or synthesis. This data suggests that C9orf72/SMCR8 complex regulates both the phosphorylation state and subcellular distribution of RAB8A.

**Figure 7.**
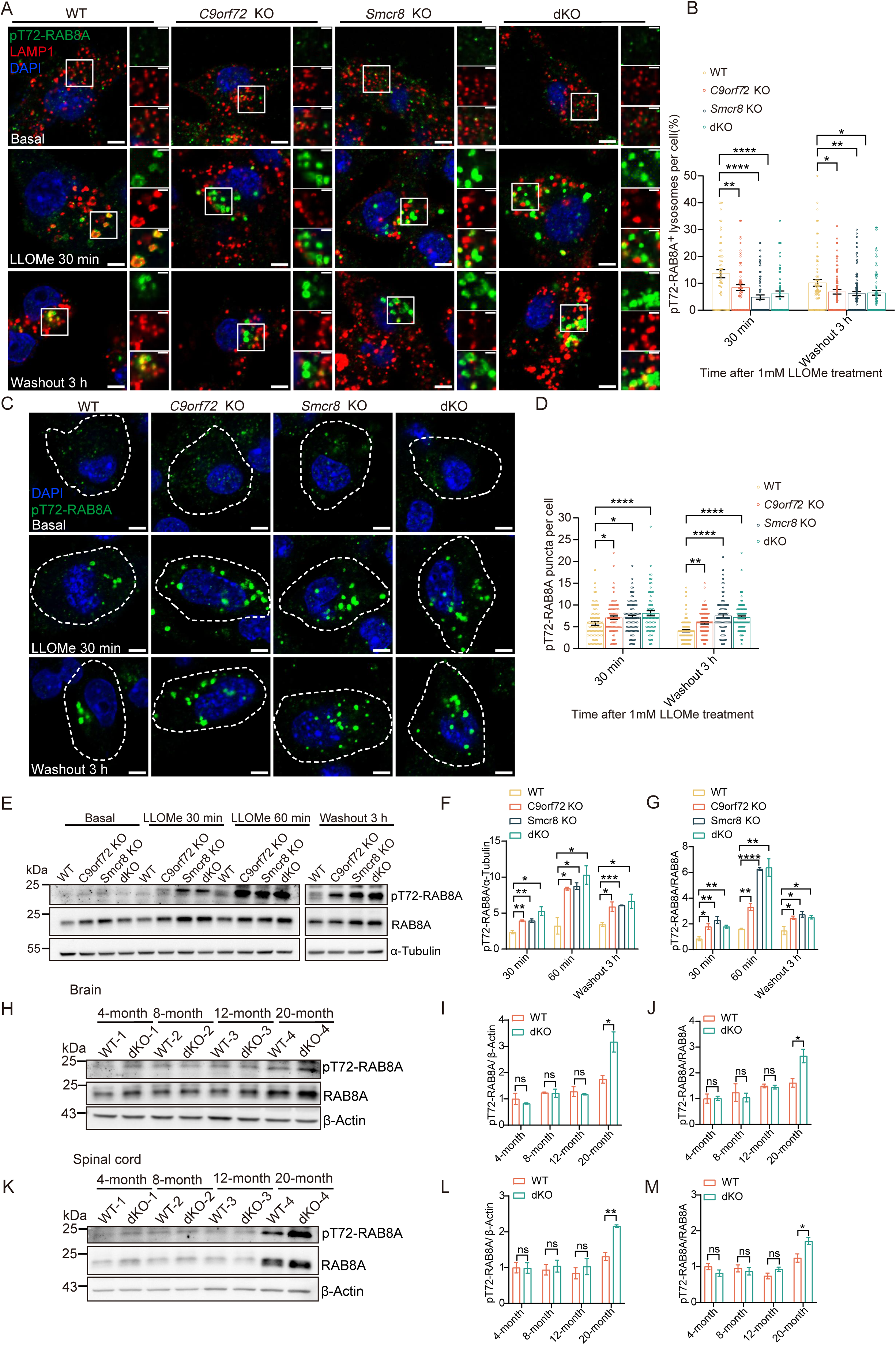
*C9orf72/Smcr8* deficiency causes RAB8A hyperphosphorylation and impaired lysosomal recruitment after lysosomal damage. (A) Representative immunofluorescence images of pT72-RAB8A (green), LAMP1 (red), and DAPI (blue) in WT and knockout BV2 cells under basal conditions, after LLOMe treatment (1 mM, 30 min), and following 3 h washout. Scale bars: 5 μm (overview), 2 μm (insets). (B) Quantification of pT72-RAB8A recruitment to lysosomes, expressed as percentage of LAMP1-positive lysosomes showing pT72-RAB8A colocalization per cell. Each dot represents an individual cell. (C) Representative immunofluorescence images of pT72-RAB8A (green) and DAPI (blue) in WT, *C9orf72* KO, *Smcr8* KO, and dKO BV2 cells under basal conditions, after 30 min LLOMe treatment, and following 3 h washout. Scale bar: 5 μm.. (D) Quantification of pT72-RAB8A-positive puncta per cell across genotypes and conditions. Each dot represents an individual cell. (E) Western blot analysis of pT72-RAB8A levels in BV2 cells (WT, *C9orf72* KO, *Smcr8* KO, and dKO) under basal conditions and at indicated time points after LLOMe treatment and washout. Blots were probed for pT72-RAB8A (top), total RAB8A (middle), and α-tubulin as loading control (bottom). (F, G) Quantification of pT72-RAB8A protein levels normalized to α-tubulin (F) or total RAB8A expression (G). (H, K) Western blot analysis of pT72-RAB8A levels in brain (H) and spinal cord (K) tissues from WT and dKO mice at 4, 8, 12, and 20 months of age. Tissue lysates were probed for pT72-RAB8A (top), total RAB8A (middle), and β-actin as loading control (bottom). (I, J, L, M) Quantification of pT72-RAB8A protein levels normalized to β-actin (I, L)or total RAB8A expression(J, M) in brain and spinal cord tissues. Data are presented as mean ± SEM from three independent experiments. For immunofluorescence quantification, ≥60 cells were analyzed per genotype per condition. Statistical analysis was performed using two-way ANOVA followed by Bonferroni post hoc test (B, D) and unpaired two-tailed Student’s t-test for pairwise comparisons (F, G, I, J, L, M). *P < 0.05, **P < 0.01, ***P < 0.001; ns, not significant.

### Loss of *C9orf72/Smcr8* leads to accumulation of GTP-Bound RAB8A and enhanced phosphorylation

Given previous findings that RAB8A is a substrate of the C9orf72/SMCR8 GAP complex in ciliogenesis, where RAB8A showed abnormal ciliary accumulation in C9orf72/SMCR8-deficient cells (Tang *et al*., 2023), we investigated whether this GAP relationship persists during lysosomal damage. While examining BV2 microglial cells under LLOMe treatment, we did not detect any ciliary signals with AC-tubulin antibodies, suggesting that the GAP activity might regulate different RAB8A pools in distinct cellular contexts (Figure S4C). We overexpressed GFP-tagged C9orf72 and SMCR8 in BV2 cells and performed co-immunoprecipitation following LLOMe treatment. Both SMCR8-GFP and C9orf72-GFP were found to associate with pT72-RAB8A under these conditions. Molecular dynamics simulations further unveiled that the phosphorylation of RAB8A enhanced its binding stability with the C9orf72/SMCR8 complex (Figure 8A, B). To directly assess the GAP relationship, we performed co-immunoprecipitation experiments using GFP-tagged RAB8A variants (wild-type, GTP-bound Q67L, and GDP-bound T22N) following LLOMe treatment.

**Figure 8.**
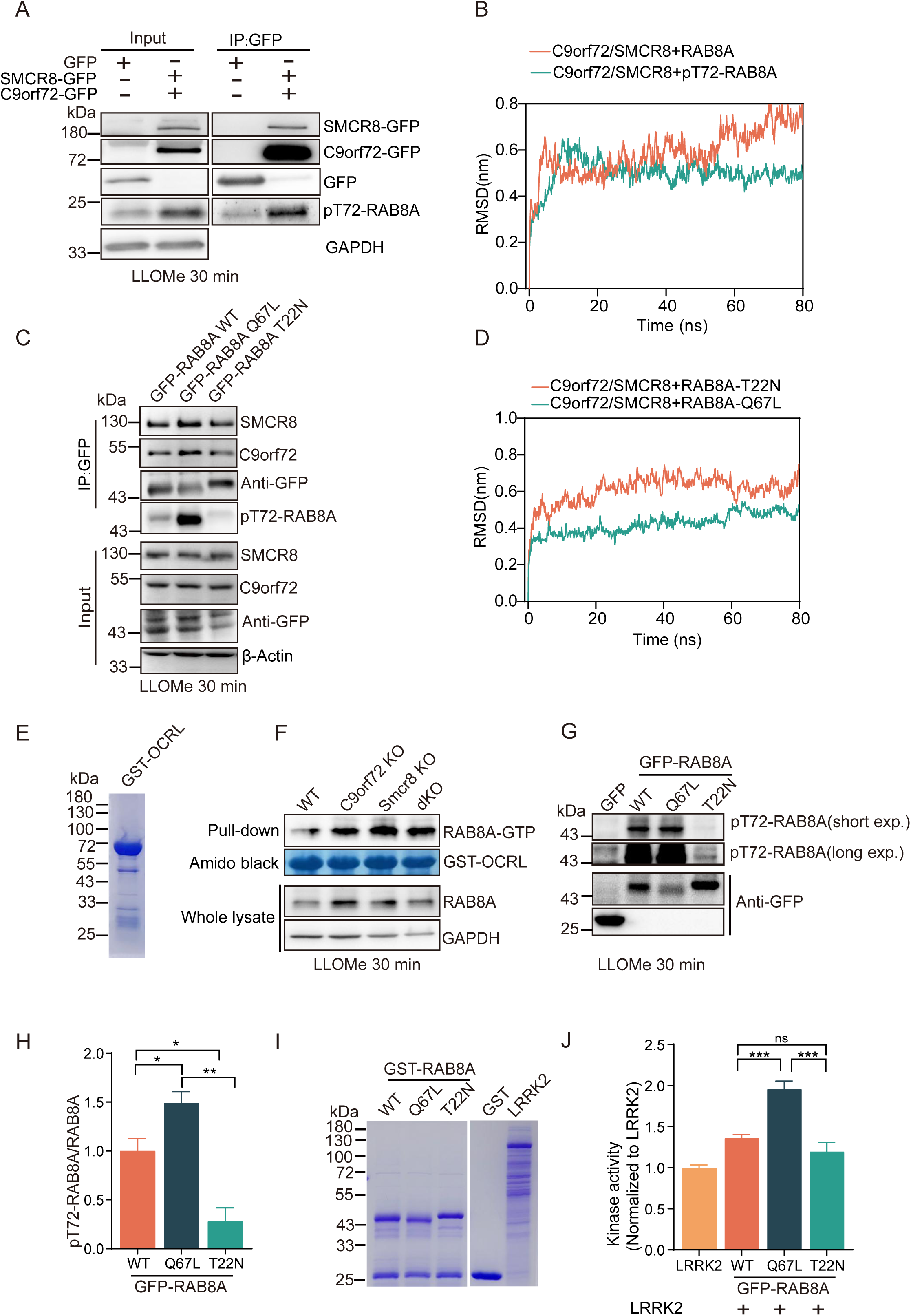
GTP-bound RAB8A shows enhanced interaction with the C9orf72/SMCR8 complex and promotes T72 phosphorylation. (A) Co-immunoprecipitation analysis of BV2 cells expressing SMCR8-GFP, C9orf72-GFP, or control vector after LLOMe treatment (1 mM, 30 min). (B) Molecular dynamics simulation showing enhanced stability of the C9orf72-SMCR8-pT72-RAB8A complex (green) compared to the C9orf72-SMCR8-RAB8A complex (orange) over simulation time. (C) Co-immunoprecipitation analysis of RAB8A variants in HEK293T cells treated with LLOMe (1 mM, 30 min). (D) Molecular dynamics simulation demonstrating greater stability of C9orf72-SMCR8 interaction with GTP-bound RAB8A Q67L (green) versus GDP-bound T22N (orange). (E, F) Coomassie blue staining of purified GST-OCRL protein (E) used as a RAB8A effector in pulldown assays, and GST-OCRL pulldown analysis (F) of lysates from WT, *C9orf72* KO, *Smcr8* KO, and dKO BV2 cells treated with LLOMe (1 mM, 30 min), followed by Western blotting for GTP-bound RAB8A. (G, H) GFP immunoprecipitation of RAB8A variants (WT, Q67L, T22N) from HEK293T cells treated with LLOMe (1 mM, 30 min), followed by Western blotting for pT72-RAB8A. Target bands are detected above 43 kDa. Quantification of pT72-RAB8A levels normalized to total RAB8A is shown in (H). (I, J) *In vitro* LRRK2 kinase assay using recombinant RAB8A protein variants. Coomassie staining confirms equal protein loading (I). Quantification (J) demonstrates enhanced kinase activity toward the GTP-bound RAB8A Q67L compared with other variants. Data are presented as mean ± SEM from three independent experiments. Statistical analysis was performed using unpaired two-tailed Student’s t-test. *P < 0.05, **P < 0.01, ***P< 0.001, ns: not significant.

Co-immunoprecipitation experiments and molecular dynamics simulations demonstrated that the C9orf72/SMCR8 complex showed preferential binding to the GTP-bound form of RAB8A (Figure 8C, D). OCRL, which specifically binds the GTP-bound form of RAB8A (Hou *et al*, 2011), was used as bait to quantify the cellular pool of GTP-bound RAB8A. We next investigated which form of RAB8A was present in mutant cells following LLOMe treatment. These results showed that the amount of the GTP-bound form of RAB8A immunoprecipitated by OCRL was increased in both single knockout and double knockout cells (Figure 8E, F). Notably, the GTP-bound (Q67L) form exhibited enhanced phospho-T72 signals, suggesting that the GTP-bound state promotes RAB8A phosphorylation (Figure 8 C, G and H). To further examine this relationship, we performed *in vitro* kinase assays using purified GST-fused RAB8A variants and LRRK2-HIS. Using an ATP-glo kit to measure phosphorylation, we confirmed that the GTP-bound form undergoes enhanced phosphorylation (Figure 8I, J). These findings establish that the increased phosphorylation of RAB8A in C9orf72/SMCR8-deficient cells is primarily due to the accumulation of its GTP-bound form, which serves as a preferred substrate for LRRK2-mediated phosphorylation.

### Rescue of lysosomal repair requires C9orf72/SMCR8 GAP activity

To establish whether the GAP activity of C9orf72/SMCR8 complex is essential for lysosomal repair, we performed rescue experiments in dKO cells. We first generated key mutations in both proteins based on previous structural and functional studies. The C9orf72 W33A mutation targets a residue in the longin domain that is critical for RAB8A substrate recognition, while the SMCR8 R147A mutation resides in the catalytic domain and has been demonstrated to abolish the GAP activity of the C9orf72-SMCR8 complex toward RAB8A(Su *et al*, 2021; Su *et al*., 2020; Tang *et al*., 2020; Tang *et al*., 2023) (Figure 9A). We then generated lentiviral constructs expressing either wild-type or mutant proteins and confirmed their expression in dKO cells by Western blot (Figure 9 B). The expression of wild-type C9orf72/SMCR8 complex in dKO cells effectively rescued the accumulation of damaged lysosomes, as evidenced by reduced Gal3-positive lysosomes after 6 hours of LLOMe washout (Figure 9C, D). In contrast, GAP-deficient mutants (C9orf72 W33A and SMCR8 R147A) failed to rescue these defects, with Gal3-positive lysosome levels remaining as elevated as in the untransfected dKO controls. These results demonstrate that both proper RAB8A binding and GAP activity of the C9orf72/SMCR8 complex are essential for efficient lysosomal repair.

**Figure 9.**
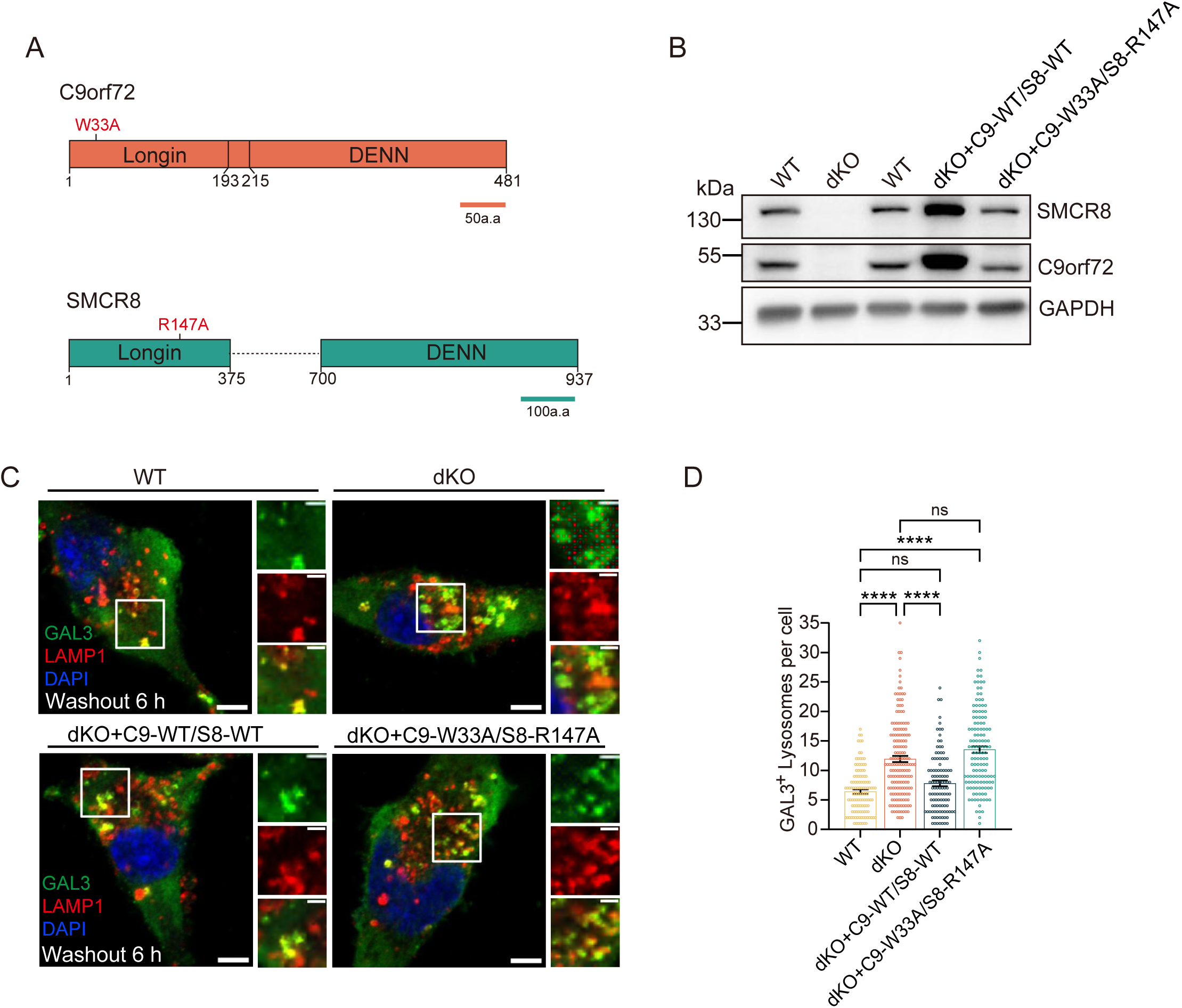
GAP activity of C9orf72/SMCR8 is required for lysosomal damage rescue. (A) Schematic representation of C9orf72 and SMCR8 protein domain architecture. C9orf72 contains a longin domain (W33A GAP-deficient mutation indicated) and a DENN domain. SMCR8 contains a longin domain (R147A GAP-deficient mutation indicated) and a DENN domain. (B)Western blot analysis showing expression levels of wild-type and GAP-mutant proteins in dKO cells. (C) Representative immunofluorescence images of dKO cells expressing WT or GAP-mutant proteins 6 h after LLOMe washout. GAL3 (green), LAMP1 (red), DAPI (blue). Scale bar: 5 μm. Insets show higher magnification of individual puncta. (D) Quantification of GAL3-positive lysosomes per cell across rescue conditions. Each dot represents one cell. Data are presented as mean ± SEM from three independent experiments. For rescue experiments, ≥110 cells were analyzed per condition across experiments. Statistical analysis was performed using one-way ANOVA followed by Tukey’s multiple comparison test. ****P < 0.0001, ns: not significant.

## Discussion

Our study identifies the C9orf72/SMCR8 complex as a central regulator of microglial homeostasis by safeguarding lysosomal repair. Loss of this complex in mice causes progressive neuroinflammation, microglial activation with DAM/MGnD-like features, and age-dependent neurodegeneration, thereby identifying impaired lysosomal repair as a key mechanism linking C9orf72/SMCR8 deficiency to microglial dysfunction (Synopsis).

The age-dependent neuroinflammatory phenotype observed in C9orf72/SMCR8-deficient mice provides insights into how chronic loss of this complex affects microglial function over time (Figure 1), consistent with the accumulation of defective lysosomes revealed in our *in vitro* and *in vivo* experiments (Figure 3,4,5 and Figure S3). Single-nucleus RNA sequencing revealed that C9orf72/SMCR8-deficient microglia adopt a disease-associated microglia (DAM)-like transcriptional profile, with upregulation of Apoe, Axl, Lgals3, and other inflammatory markers (Figure 2) (Deczkowska *et al*, 2018; Depp *et al*, 2025; Fumagalli *et al*., 2025). This phenotype mirrors disease-associated microglia (DAM) and the microglial neurodegenerative phenotype (MGnD), both of which have been implicated in neurodegenerative conditions. This state was accompanied by enhanced phagocytic activity, which may drive inappropriate synaptic pruning and neuronal loss. Such a mechanism is consistent with previous studies showing that C9orf72-deficient microglia in Alzheimer’s disease models exhibit hyperactive synapse engulfment and accelerate cognitive decline(Lall *et al*., 2021; Lui *et al*, 2016).

Our data further implicate defective lysosomal repair as the underlying driver of this phenotype. *In vivo*, we first observed that aged C9orf72/SMCR8-deficient mice exhibited elevated GAL3 levels in brain and spinal cord. Immunofluorescence revealed that these increases reflected enhanced GAL3-positive lysosomes specifically in microglia, indicating accumulation of damaged lysosomes. Consistently, *in vitro* experiments showed that C9orf72/SMCR8-deficient microglia and BV2 cells accumulated persistent GAL3-positive lysosomes following LLOMe-induced lysosomal stress, reflecting defective recovery (Figure 3). We demonstrate that the complex is recruited to injured lysosomes and facilitates ESCRT engagement, thereby promoting membrane repair (Figure S3, Figure 6). Loss of either subunit (C9orf72/SMCR8) leads to accumulation of damaged lysosomes, increased galectin-3 puncta, and impaired ESCRT recruitment (Figure 5C, Figure 6C). Thus, impaired recruitment of ESCRT components in knockout cells provides a mechanistic explanation for the persistence of GAL3-positive lysosomes. The progressive accumulation of GAL3-positive lysosomes in aged knockout brain and spinal cord suggests that impaired repair amplifies with age, as damaged lysosomes accumulate through lipid and protein overload(Choi *et al*, 2022; Gabandé-Rodríguez *et al*, 2019; Marschallinger *et al*, 2020). Importantly, we observed that single knockouts retained only minimal amounts of the partner protein (Figure S2C), with most phenotypes being comparable between single and double knockouts, consistent with previous findings that C9orf72 and SMCR8 are mutually dependent for protein stability(Shao *et al*., 2020; Zhang *et al*., 2018),

Mechanistically, we reveal a dual role for the C9orf72/SMCR8 complex in regulating RAB8A during lysosomal repair. This complex not only functions as a GAP for RAB8A but also influences its phosphorylation dynamics (Figure 8). While this complex was previously shown to functions as a RAB8A GAP ciliogenesis (Tang *et al*., 2023), we demonstrate that it regulates RAB8A in a cilium-independent context in microglia during lysosomal stress. Notably, we did not detect primary cilium formation in BV2 cells following LLOMe treatment (Figure S4C), consistent with the non-ciliated nature of microglial cells(Sipos *et al*, 2018; Sterpka & Chen, 2018). Mutant cells accumulated GTP-bound RAB8A, which is preferentially phosphorylated by LRRK2(Figure 8). Consequently, both single and double knockouts displayed elevated phospho-RAB8A levels *in vitro* and in aged CNS tissue (Figure 7). Strikingly, however, phospho-RAB8A failed to localize to damaged lysosomes and instead accumulated in cytoplasmic puncta, thereby preventing ESCRT recruitment (Figure 7). This mislocalization distinguishes the C9orf72/SMCR8-deficient state from Parkinson’s disease-associated LRRK2 mutations, which cause excessive RAB8A phosphorylation and sequestration at lysosomes and centrosomes(Madero-Pérez *et al*, 2018; Mamais *et al*, 2021). Thus, C9orf72/SMCR8 deficiency produces a unique defect: phosphorylation of RAB8A is increased, its spatiotemporal targeting to damaged lysosomes is impaired. Rescue experiments with wild-type, but not GAP-deficient, C9orf72/SMCR8 mutants confirmed that the GAP activity of this complex is indispensable for lysosomal repair (Figure 9).

In conclusion, our findings establish impaired lysosomal repair as a key mechanism linking C9orf72/SMCR8 deficiency to microglial dysfunction, thereby providing a mechanistic framework that may explain neuroinflammation in ALS and FTD. By uncovering disrupted ESCRT recruitment and aberrant RAB8A dynamics, we provide a mechanistic framework that extends the functional repertoire of the C9orf72/SMCR8 complex beyond its known roles. These findings advance our understanding of C9orf72/SMCR8 function in microglia and suggest that targeting lysosomal repair pathways or RAB8A regulation could represent potential therapeutic strategies for conditions involving microglial dysfunction.

Limitation of this study

This study primarily utilized mouse models, and it would be particularly meaningful to investigate whether ALS/FTD patients exhibit increased lysosomal damage, especially in microglial cells. Our discovery of hyperphosphorylated RAB8A in mutant cells represents a potential biomarker for ALS/FTD that warrants validation in patient tissues or iPSC-derived microglia. Additionally, whether pharmacological or genetic inhibition of RAB8A hyperphosphorylation can mitigate lysosomal repair defects remains to be determined. Since we did not observe changes in RAB10 phosphorylation status, another damage repair protein, further studies are needed to determine RAB10’s potential involvement in this process. Lysosomal damage repair also involves lipid transfer from the endoplasmic reticulum to facilitate damaged lysosome repair. Our current antibody limitations prevent adequate detection of ER-lysosome contacts, representing an important future research direction. Understanding these mechanisms may provide therapeutic targets for C9orf72-related neurodegeneration.

## Materials and Methods

### Animals

*C9orf72* knockout mice (Jackson Laboratory, stock #027068) and *Smcr8* knockout mice (as described in Shao *et al.,* 2020) were maintained on a C57BL/6J background. Double knockout mice were generated by intercrossing *C9orf72^-/-^* and *Smcr8^-/-^* lines. All mice were housed in pathogen-free barrier facilities under a 12-hour light/dark cycle at 21 ± 2 °C and 40–60% humidity. All experimental procedures were approved by the Institutional Animal Care and Use Committee of Yunnan University. Age-matched wild-type, heterozygous and double knockout littermates ranging from 3 to 24 months were used for experimental analyses.

### snRNA-seq samples and data processing

Single-nucleus RNA sequencing (snRNA-seq) was performed on pooled brain samples from 20- to 24-month-old mice, with three independent WT brains combined and three independent dKO brains combined to generate the sequencing libraries. The raw data were aligned to the mouse genome GRCm39 using DNBC4TOOLS (v2.1.3). The data analysis and visualization were carried out using the Seurat R package (v5.0)(Hao *et al*, 2024). Cells were filtered based on the following criteria: 100 < nFeatures < 7500 and 200 < nCountFeatures< 30000. The cells with percent.mt > 20 were further excluded. The data were normalized by using NormalizeData function with default parameters. The data was then scaled by using ScaleData function with vars.to.regress = c (“percent.mt”, “percent.Rpsl”, “percent.Hb”. After that, the data was integrated by using IntegrateLayers function with method = HarmonyIntegration. Unsupervised clustering was then performed by FindNeighbors function with dims=1:30 and FindClusters function with resolution = 0.05, and the cells was annotated by using the published mouse brain transcriptomic atlas(Yao *et al*., 2021). A total of 33,257 cells were obtained, including 14,460 excitatory neurons, 9,379 inhibitory neurons, 6,267 oligodendrocytes, 1,912 microglia, 832 astrocytes, and 407 epithelial cells. Differentially expressed genes (DEGs) between KO and WT conditions within each cell-type were identified using the FindMarkers function with default parameters. Functional enrichment analysis of Kyoto Encyclopedia of Genes and Genomes (KEGG)(Ogata *et al*, 1999) and Gene Ontology (GO)(Ashburner *et al*, 2000) were performed using the clusterProfiler package (v4.10.1)(Wu *et al*, 2021; Yu *et al*, 2010; Yu *et al*, 2012). In brief, the enrichGO function was utilized to identify enriched GO terms with (ont= “BP”/ “MF”/ “CC”, pvalueCutoff =0.05, qvalueCutoff = 0.2) and then the similar GO terms were merged based on semantic similarity using the simplify function with cutoff=0.7. The KEGG enrichment analysis was performed using enrichKEGG function with (pvalueCutoff =0.05, qvalueCutoff =0.2, minGSSize=5).

### Immunostaining of sections and cultured cells

Mice aged 4 to 24 months were anesthetized with CO₂ and transcardially perfused with ice-cold PBS, followed by perfusion-fixation with 4% paraformaldehyde (PFA) in PBS. Brains were collected and post-fixed in 4% PFA at 4 °C for 24 h. After washing with PBS, tissues were cryoprotected in 30% sucrose for 2 days and then embedded in OCT. Coronal sections (15 µm) were prepared using an HM525 NX cryostat (Thermo Fisher Scientific). Sections were washed three times in PBS (5 min each) and incubated in blocking buffer (2% normal goat serum, 1% BSA, 0.1% Triton X-100 in PBS) for 1 h at room temperature. Subsequently, sections were incubated with primary antibodies diluted in blocking buffer overnight at 4 °C. After three PBS washes, sections were incubated with secondary antibodies in blocking buffer for 1 h at room temperature. Nuclei were counterstained with DAPI, and sections were mounted with antifade mounting medium. Once dried, slides were stored at 4 °C until imaging.

For immunostaining of cultured cells, coverslip-seeded cells were fixed with freshly prepared 4% PFA for 10 min at room temperature and washed three times with PBS. Cells were permeabilized with 0.1% Tween-20 for 10 min, washed, and then blocked with 2% normal goat serum and 1% BSA in PBS for 1 h at room temperature. Cells were incubated with primary antibodies in blocking buffer overnight at 4 °C, washed three times with PBS, and incubated with secondary antibodies in blocking buffer for 45 min at room temperature. After final PBS washes, coverslips were mounted in antifade medium. Images were acquired using a Zeiss LSM 800 confocal laser scanning microscope.

### Cells, Transfection, and Reagents

BV2 cells, a gift from Dr. Q.L., and 293T cells were cultured at 37°C with 5% CO2 in DMEM supplemented with 10% FBS (Vivacell), 100 U/ml penicillin, and 100 mg/ml streptomycin. Transfections for BV2 cells were performed using the Glial-Mag kit (Oz Biosciences) according to the manufacturer’s instructions. Briefly, 1×10^5^ BV2 cells were plated on an 18mm coverslip in a 24-well plate, with a ratio of 2.5 μl Glial-Mag to 1 μg DNA. 250 μl Glial-Boost was added to the well and incubated for 1 hour. The transfection mix was then removed, and imaging was captured, or immunostaining was performed after 24 hours. Transfections for 293T cells were performed using Polyethylenimine ((PEI 40K, MX2203)), with a ratio of 4 μl PEI to 1 μg DNA.

### Generation of *c9orf72*, *smcr8* single and double KO cell lines

The CRISPR/Cas9 technology was used to knock out C9orf72 and Smcr8 genes in mouse BV2 cell line. First, the complete sequence information of mouse C9orf72 and Smcr8 genes was obtained from the NCBI website to determine the sgRNA targeting sequences. The first exon was chosen as the target fragment for designing sgRNA to ensure that only the long isoform of the C9orf72 gene and all isoforms of the Smcr8 gene were knocked out. The sgRNA sequences for C9orf72 (C9-sgRNA-2F: CACCGCACACACTCTGTGAAGTGGG, C9-sgRNA-2R: AAACCCCACTTCACAGAGTGTGTGC) and Smcr8 (S8-sgRNA-2F: CACCGTGATGTGGTGGCCTTCACCA, S8-sgRNA-2R: AAACTGGTGAAGGCCACCACATCAC) were designed using the website https://portals.broadinstitute.org/gppx/crispick/public and synthesized with sticky ends for ligation reactions and constructed into lentiCRISPRv2 vectors, followed by lentivirus packaging. For virus infection, BV2 cells were seeded in a 6-well plate and infected with the virus. After puromycin selection, the surviving cells were passaged to adjust their condition and further selected with puromycin. A portion of the cells was used to detect the gene knockout efficiency by Western Blot. The cells with better knockout efficiency were sorted into single cells using flow cytometry for further experiments. The knockout of C9orf72 and Smcr8 genes in the selected single-cell clones was confirmed by Western Blot and DNA sequencing. To generate C9orf72 and Smcr8 double knockout cells, the Smcr8 knockout cells were infected with the C9orf72 sgRNA-containing plasmid, followed by single-cell selection, and the double knockout was confirmed by Western Blot and genomic DNA sequencing.

### Lyso-IP Immunoblot

Lyso IP immunoblot assay was performed according to previously described protocols.(Abu-Remaileh *et al*, 2017). To establish Lyso IP stably expressing cell line, the pLenti-TMEM192-3×HA-Blasticidin construct was packaged into lentiviral particles combination with pCMV-VSVG and psPAX2 packaging plasmids in HEK293T cells. BV2 C9orf72 knockout and Smcr8 knockout cells were seeded in 6 cm culture dish and infected with 700 µL of TMEM192-3xHA virus. 24 h post-infection, 6 µg/mL Blasticidin S HCl was added for selection. Following expansion and counting, 8.0–8.5×10⁶ cells were seeded in 10 cm culture dishes. After cells reached ∼85% confluency. Cells were washed twice with pre-cold 1× PBS and once with pre-cold KPBS buffer (136 mM KCl,10 mM KH₂PO₄, pH 7.25 adjusted with KOH). Subsequently, 500 µL KPBS supplemented with protease and phosphatase inhibitors was added, and cells were gently scraped on ice. Cell suspensions were gently homogenized for 120 strokes use a 2 mL Dounce homogenizer (B pestle) (D8938, Sigma-Aldrich) until ∼ 80% trypan blue-positive cells were observed. The lysate was centrifuged at 3000 rpm for 2 min at 4°C to remove pellet nuclei and cellular debris. The supernatant was incubated with 80 µL KPBS prewashed anti-HA magnetic beads (88836, Thermo Fisher) at 4°C for 15 min with gentle rotation. Beads were washed five times with KPBS, then resuspended in 50 µL of 1× Laemmli sample buffer, and boiled at 95 °C for 10 min. Eluted proteins were analyzed by Western blotting.

### Protein extraction and immunoblotting analysis

Cells were collected by centrifugation at 300 g for 4 min at 4°C and washed twice with cold PBS. Cell lysates for western blot were prepared using protein lysis buffer (50 mM Tris-HCl, pH 7.5, 150 mM NaCl, 1% Triton X-100, 1 mM EDTA) supplemented with protease inhibitor cocktail (MCE) and incubated on ice for 30 min. After centrifugation at 17,000 g for 20 min at 4°C, cleared supernatant was transferred to new clean tubes, and total protein concentration was measured using BCA Protein Assay kit (Pierce).

Protein samples were boiled for 10 min at 95°C in 1xSDS sample buffer with β-mercaptoethanol (1610747, BioRad), resolved on 10% polyacrylamide gels or pre-cast 4–15% Mini-PROTEAN® TGX™ Precast Protein Gels (#4561094), and transferred to a PVDF membrane (IPFL00010; Millipore) using a wet transfer module (BIO-RAD Mini-PROTEAN Tetra System). Membranes were blocked in 5% non-fat milk powder, and resuspended in PBS-T for 30min at room temperature, followed by incubation with primary antibodies diluted in PBST 5% milk overnight at 4°C. After washing with PBST, membranes were developed using Ultra High Sensitivity ECL Kit (BL520B) Western blotting images were acquired by SINSAGE smartchemi 610 and the grayscale value of protein bands was quantified using ImageJ software.

### Immunoprecipitation and GST pull-down

For GFP-based IP, BV2 cells expressing GFP-tagged C9orf72 and SMCR8 were treated with 1 mM LLOMe for 30 min. Cells were washed three times with PBS, and lysed in lysis buffer containing 20 mM Tris-HCl, pH 7.4, 150 mM NaCl, 0.5 % NP-40, Protease Inhibitor Cocktail and Phosphatase Inhibitor Cocktail. After unbroken cells and large debris were removed, the supernatant was incubated with GFP-trap beads (ChromoTek) overnight at 4°C. The incubated beads were washed five times with the wash buffer containing 20 mM Tris-HCl, pH 7.4, 150 mM NaCl, 0.5% NP-40, 1 mM PMSF. Immunoprecipitated proteins were evaluated with western blotting.

For GST pull-down, BV2 wild-type (WT), C9orf72 knockout (KO), Smcr8 knockout (KO) and double knockout (dKO) cell lysate were incubated with GST-OCRL proteins immobilized on glutathione Sepharose beads at 4°C overnight. The beads were washed five times with the same wash buffer and then analyzed by western blotting.

### Kinase activity assay

The kinase activity assay using a ADP-Glo™ Kinase Assay kit (Promega, V6930) according to the manufacturer’s protocol. 5 μg recombinant LRRK2 protein, were incubated with 9 μg substrates [WT-RAB8A, GTP-bound RAB8A (Q67L) and GDP-bound (T22N)] in reaction buffer. After in vitro kinase assay, ADP-Glo reagent and kinase detection reagent was added to terminate the reaction and deplete residual ATP. The generated ADP is subsequently converted to ATP, and the newly formed ATP is quantified by chemiluminescence.

### Transmission Electron Microscopy

BV2 cells were grown in 24-well plates in DMEM. Cells were fixed with PBS containing 2.5% glutaraldehyde (02607-BA, SPI Supplies) at 4°C and washed with PBS. Samples were post-fixed in 1% osmium tetroxide (OsO₄, 02602-AB, SPI Supplies) in distilled water at 4°C and subsequently rinsed with distilled water. En bloc staining was performed at room temperature with 2% aqueous uranyl acetate (02624-AB, SPI Supplies), followed by distilled water washes. Samples were dehydrated through a graded ethanol series, then infiltrated using a graded series of EMBED-812 resin diluted in anhydrous ethanol (EMBED-812, 14900, NMA, 19000, DDSA, 13710, Electron Microscopy Sciences). After overnight infiltration in 100% resin at room temperature, samples were further infiltrated with 100% resin containing the accelerator DMP-30 (13600, Electron Microscopy Sciences), embedded in molds, and polymerized at 60°C. Ultrathin sections (∼70 nm) were cut on an ultramicrotome (EM UC7, Leica Biosystems) and collected on copper grids. Sections were post-stained with 2% aqueous uranyl acetate and then with 0.2% lead citrate (A10701, Alfa Aesar), rinsed with distilled water and air-dried. Specimens were examined using a transmission electron microscope (HT7800, Hitachi) operated at an accelerating voltage of 80 kV.

### Primary microglia isolation

Primary microglia were isolated following the procedure described previously(Du *et al*, 2021). Postnatal mice were anesthetized by placing them on ice for 30-50 s. After sterilization with 75% ethanol, all pups were quickly decapitated with surgical scissors and put in the ice-cold 1x PBS. Meanwhile, rinse all heads with ice-cold 1x PBS to remove excess blood. Transfer the heads to a new plate containing 2 mL of pre-cold 1x PBS. The olfactory bulbs and cerebellum were discarded, and the remaining brain tissue was gently transferred to a chilled 6-well plate containing 0.5 mL of pre-cold 1x PBS. The brain tissue was finely minced into 1-2 mm³ pieces with the help of spring scissors. Tissues were incubated in digestion buffer (DMEM containing 8 U/mL papain [P5306, Sigma] and 125 U/mL DNase I [DN25, Sigma]) at a ratio of 0.5 mL digestion buffer per brain tissue, at 37 °C for 20 min with gentle agitation every 5 min. The papain digestion was terminated by adding 3 mL of pre-warmed complete culture medium (DMEM/F12 supplemented with 10% fetal bovine serum and 1% penicillin-streptomycin). The digested tissue was gently triturated 10 times using a 1mL pipette. Let it settle for 1 min. Carefully pass the cell suspension through a pre-wetted 70 µm cell strainer and collect the flow-through in the 50 mL collection tube. Repeat this step until all the big clumps have been removed. The cell suspension was centrifuged at 200 ×g for 10 min at room temperature, and then discard the remaining cell debris. Primary cells were resuspended in complete culture medium supplemented with 2 ng/mL GM-CSF(HY-P7361, MCE) and seeded in culture dishes pre-coated with 0.1 mg/mL poly-D-lysine (P2636, SIGMA). On the second day, wash the dishes with pre-warmed PBS to remove non-adherent cells and debris. Fresh medium containing 2 ng/mL GM-CSF was added, and cells were maintained for 4–7 days before microglia were harvested for the desired functional assay.

### Statistics

Data derived from different genetic backgrounds or different treatments were analyzed using Student’s two-tailed unpaired t-test, one-way ANOVA followed by Tukey’s multiple comparison test or two-way ANOVA followed by Bonferroni post-test for multiple comparisons. Data are represented as mean ± standard error of the mean (SEM). A value of p < 0.05 was considered significant (*p < 0.05; **p < 0.01; ***p < 0.001).

## Abbreviations

(ALS): Amyotrophic lateral sclerosis
(FTD): Frontotemporal dementia
(GAP): GTPase-activating protein
(CNS): Central nervous system

## Acknowledgments

This research was supported by National Science Foundation of China (32070743 to M. Y., 32170662 to C.P.); Yunnan Fundamental Research Projects (202401BF070001-015 to M.Y., 202401AS070131 to C.P.); Research Startup Funding of Yunnan University(C1762201000139); The Xingdian Talents Support Program of Yunnan Province (C619300A142). The 16th Graduate Research and Innovation Project (KC-242410334). We are grateful to Dr. Jianfu Chen for providing *Smcr8* mutant mice. We also thank Dr. Qun Lu. for generously providing BV2 cells.

## Author Contributions

M.Y. and Y.W. conceived the project. S.L. performed most experiments. S.X. performed TEM analysis and assisted mouse experiments. F.L., Q.Z., X.S., Q.G., P.Z. and J.B. assisted with imaging, cell line construction, vector construction, and protein purification. H.X., and Q.C. assisted with protein purification and complex analysis. Y. T. and C. P. assisted with bioinformatics analysis. M.Y., Y.W., S.L. and S.X. wrote the manuscript, and all the authors read and commented on the manuscript.

---

**Figure S1.**
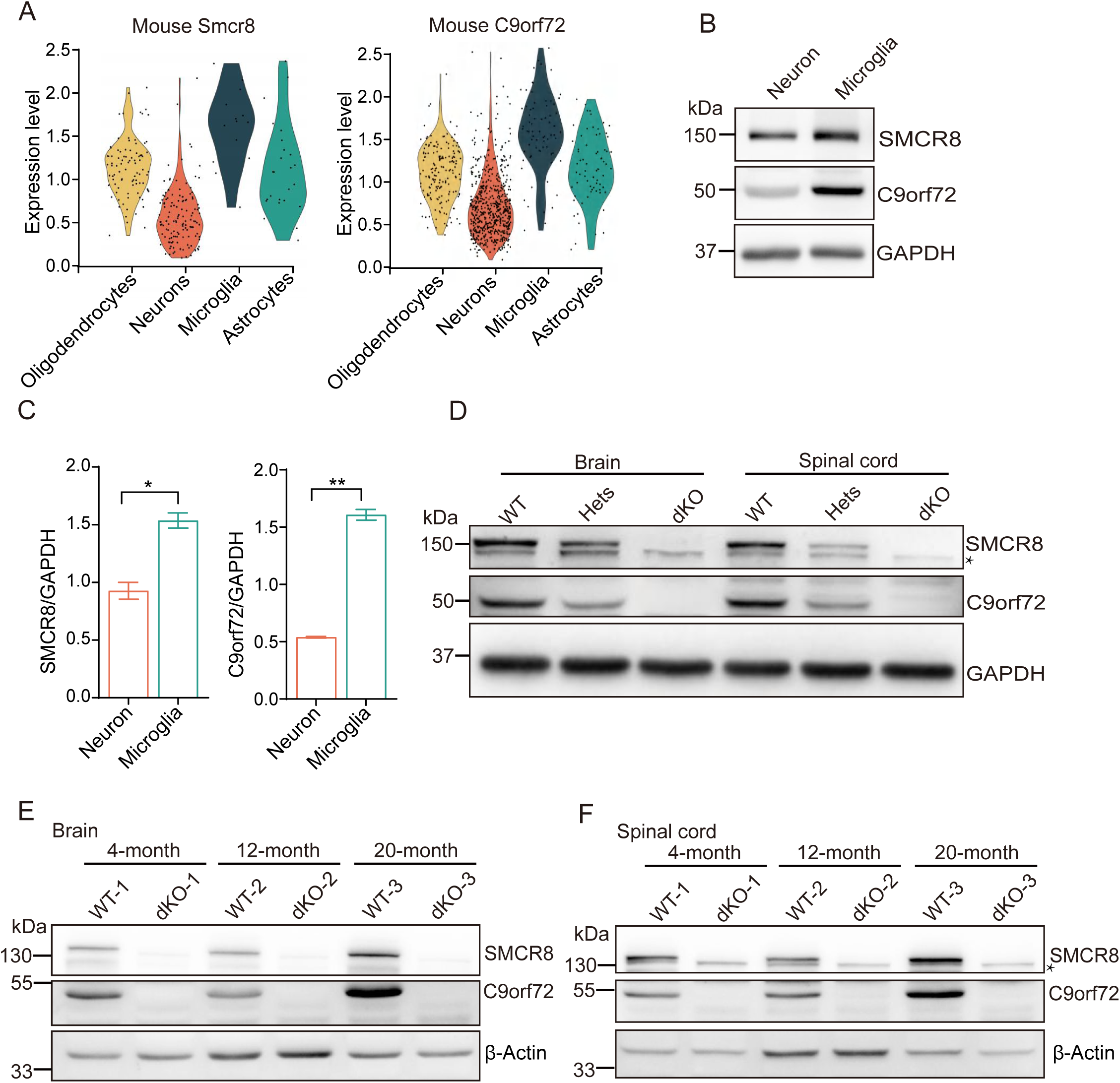
C9orf72 and SMCR8 are highly expressed in microglia and exhibit age-dependent upregulation in the CNS. (A) Violin plots showing *C9orf72* and *Smcr8* expression levels across major CNS cell types from single-cell RNA-sequencing dataset (Yao *et al*., 2021). (B, C) Western blot analysis of C9orf72 and SMCR8 in primary neurons and microglia, with GAPDH as loading control (B), and quantification of protein levels normalized to GAPDH (C). (D) Validation of knockout efficiency in C9orf72/SMCR8 double knockout (dKO) mice by Western blot analysis of brain and spinal cord tissue lysates. Representative blots show protein expression in WT, heterozygous (Het), and dKO genotypes. GAPDH serves as loading control. Asterisks indicate nonspecific bands. (E, F) Age-dependent increase of SMCR8 and C9orf72 protein levels in brain (E) and spinal cord (F) from WT mice at 4–20 months. β-Actin serves as the loading control. Asterisks indicate nonspecific bands. Data are presented as mean ± SEM (n = 3 mice per group from three independent experiments). Statistical analyses were performed using unpaired two-tailed Student’s t-test. *P < 0.05, **P < 0.01.

**Figure S2.**
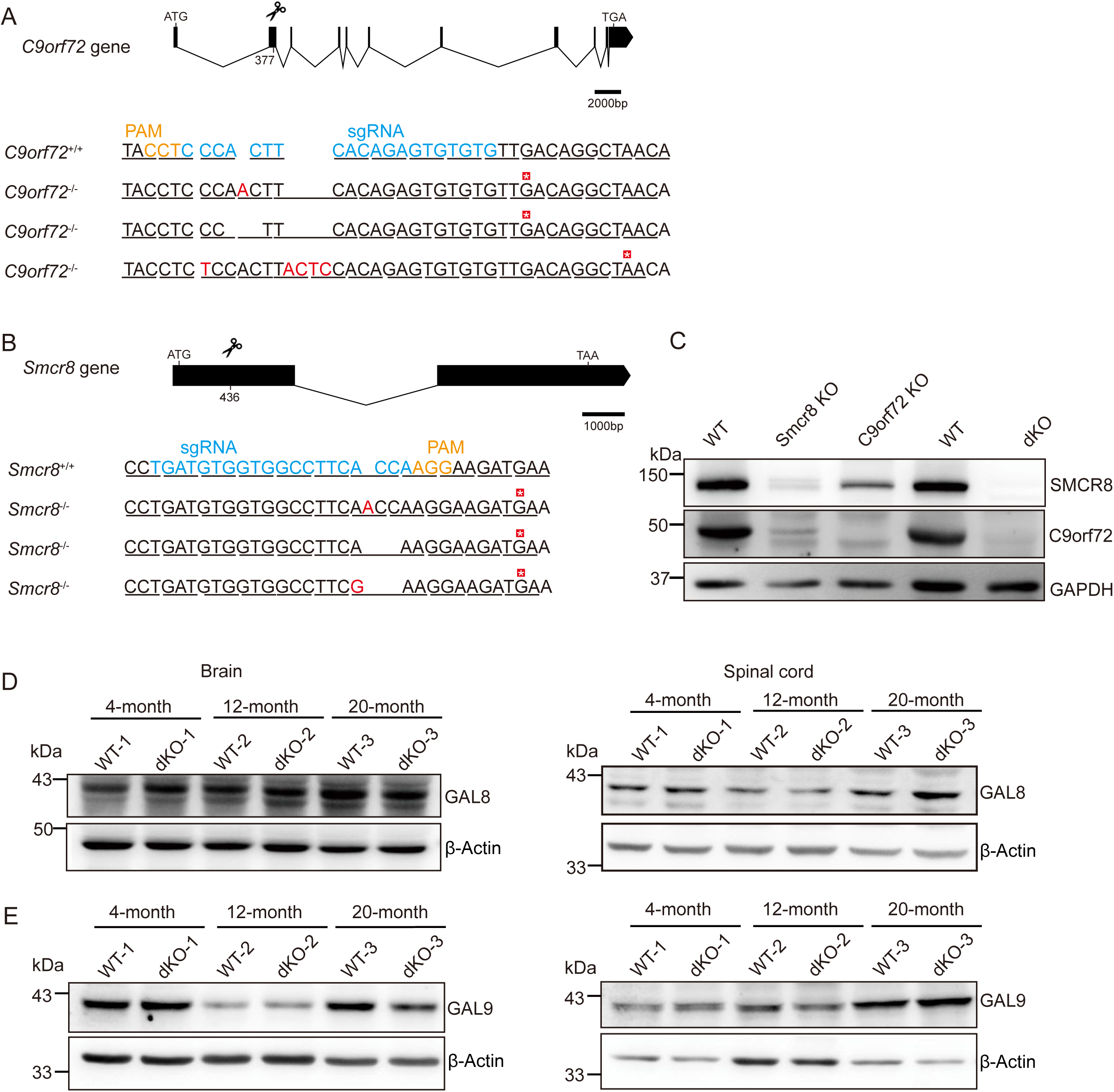
Generation and validation of *C9orf72* and *Smcr8* knockout BV2 cells. (A) Schematic of *C9orf72* gene structure and CRISPR/Cas9 targeting strategy. Representative sequencing results from a knockout clone show insertions/deletions at the target site leading to frameshift mutations and premature stop codons. Blue indicates sgRNA target sites, orange indicates PAM motifs, and asterisks mark stop codons. (B) Schematic of *Smcr8* gene structure and CRISPR/Cas9 targeting strategy. Representative sequencing results from a knockout clone show deletions causing premature termination. Double knockout (dKO) was generated by sequential targeting of *Smcr8* KO cells with *C9orf72* sgRNA. (C) Western blot analysis confirming knockout efficiency in generated cell lines. SMCR8, C9orf72, and GAPDH protein levels were assessed in WT, *Smcr8* KO, *C9orf72* KO, and dKO BV2 cells. Complete loss of target proteins confirms successful knockout generation. GAPDH serves as loading control. (D, E) Western blot analysis of GAL8 and GAL9 protein levels in brain and spinal cord tissues from WT and dKO mice. Data are representative of three independent experiments.

**Figure S3.**
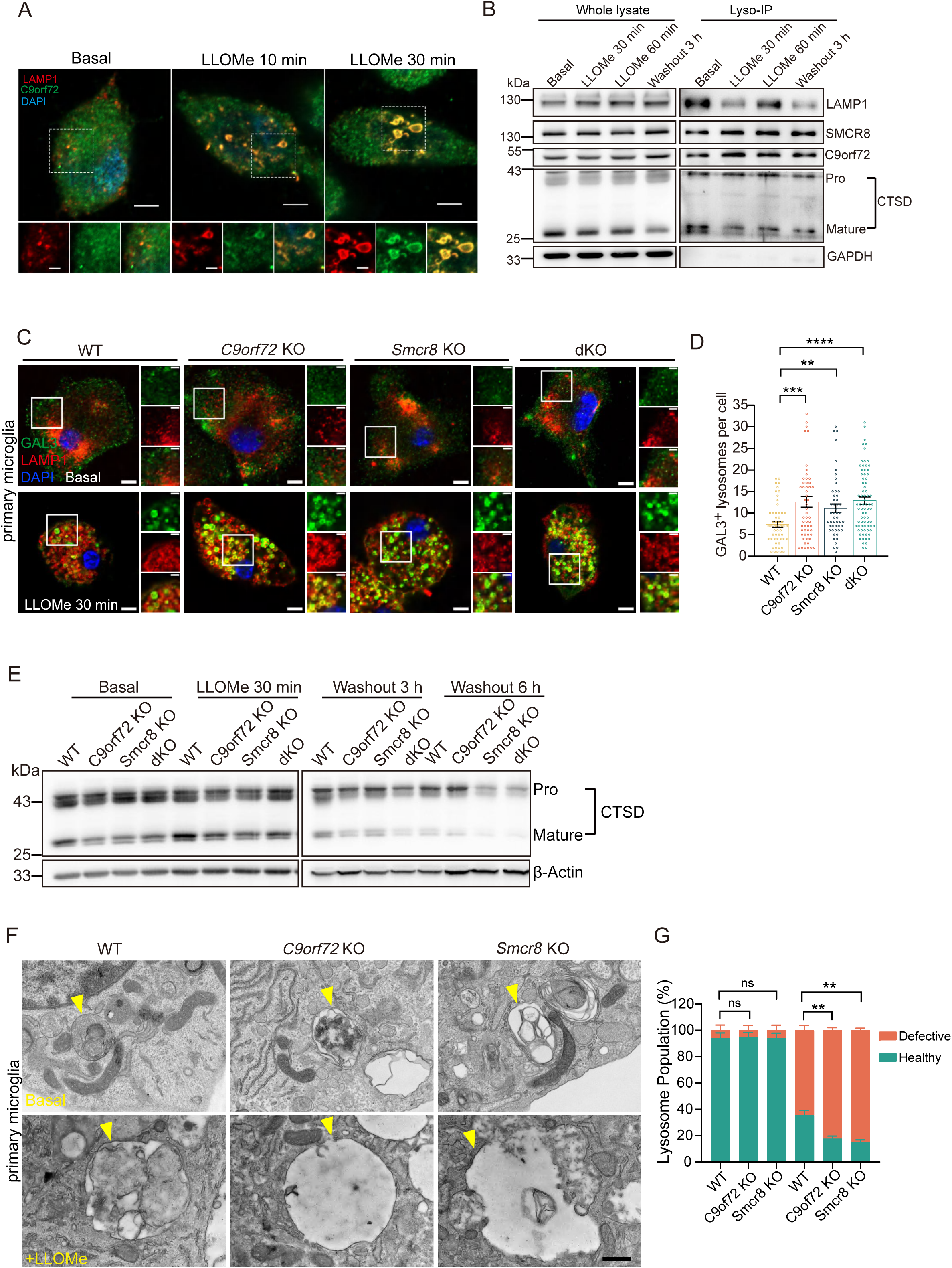
C9orf72 and SMCR8 are critical for maintaining lysosomal membrane integrity and repair. (A) Representative immunofluorescence images of endogenous C9orf72 (green) and LAMP1 (red) in microglia under basal conditions and after LLOMe treatment (10, 30 min). Colocalization regions shown in insets. Scale bars: 5 μm (overview), 2 μm (insets). (B) LysoIP analysis showing association of endogenous C9orf72 and SMCR8 with lysosomes after LLOMe treatment (1mM). Input and immunoprecipitated fractions analyzed by Western blot. (C) Representative immunofluorescence images of GAL3(green), LAMP1(red), DAPI (blue) in WT, *C9orf72* KO, *Smcr8* KO and dKO primary microglia under basal condition and after LLOMe treatment (0.5 mM, 30 min). Scale bar: 5 μm. (D) Quantification of GAL3-positive lysosomes per cell in primary microglia. Each dot represents the number of GAL3/LAMP1 double-positive structures counted in an individual cell. (E) Western blot analysis of cathepsin D (CTSD) processing under basal conditions, after LLOMe treatment (1 mM, 30 min), and during washout periods (3, 6 h). (F) Transmission electron microscopy images of primary microglia from WT, *C9orf72* KO, and *Smcr8* KO mice. Yellow arrowheads indicate defective lysosomes characterized by altered morphology, membrane disruption, and reduced electron-dense content. Scale bar, 500 nm. (G) Quantification of defective lysosomes as percentage of total lysosomal structures per cell. Defective lysosomes were identified by morphological criteria including lysosome enlargement, reduced electron density, or membrane disruption. Data are presented as mean ± SEM from three independent experiments. For immunofluorescence quantification, ≥60 cells were analyzed per genotype per condition (≥20 cells per experiment). For electron microscopy analysis, ≥20 cells were examined per genotype across three independent preparations Statistical analysis was performed using unpaired two-tailed Student’s t-test (D, G). **P < 0.01, ***< 0.001,****P < 0.0001, ns: not significant.

**Figure S4.**
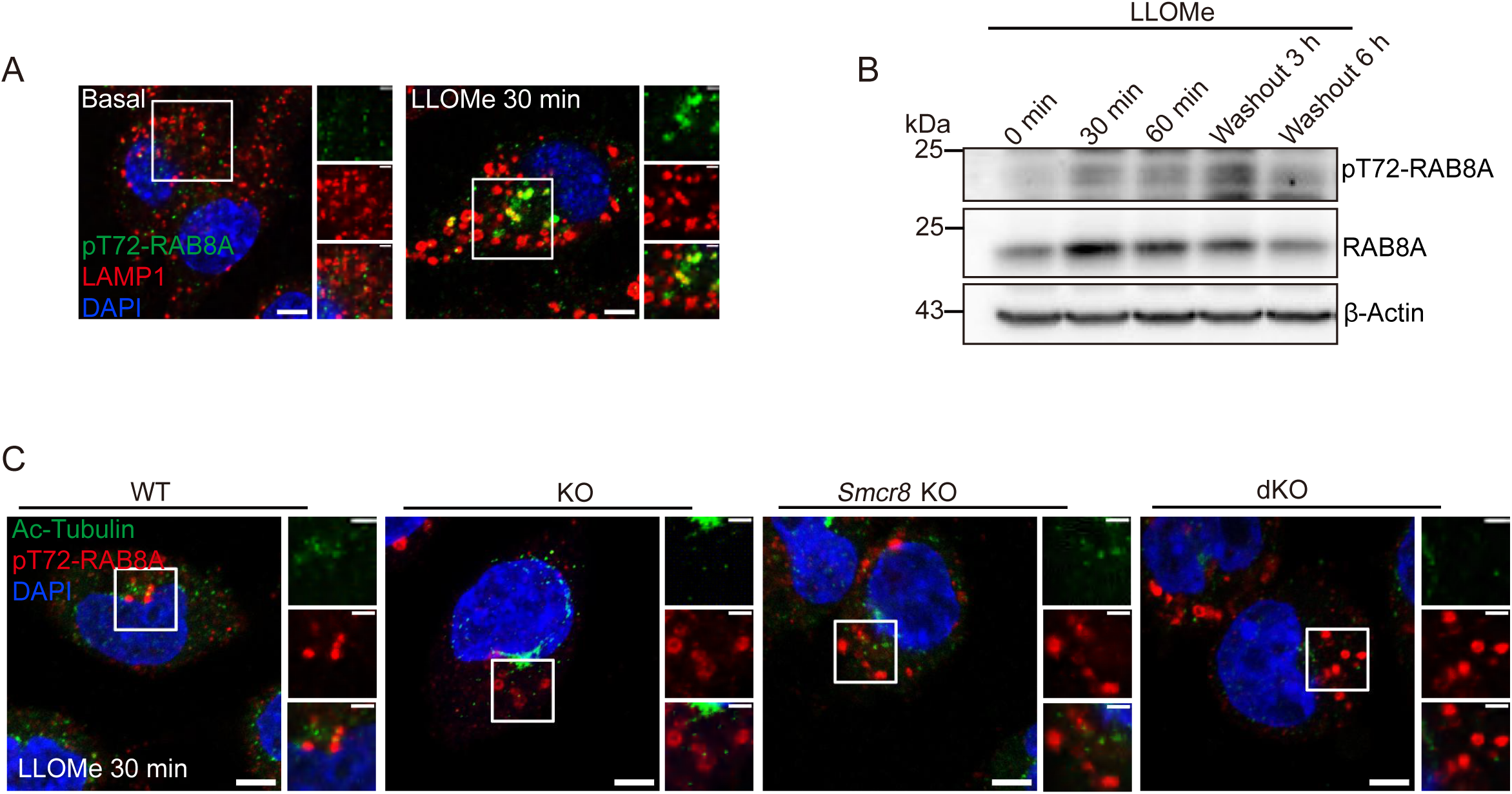
Localization and dynamics of pT72-RAB8A in BV2 cells during lysosomal damage. (A) Representative immunofluorescence images of pT72-RAB8A (green), LAMP1 (red), and DAPI (blue) in BV2 cells under basal conditions and after LLOMe treatment (1mM, 30 min). Scale bars: 5 μm (overview), 2 μm (insets). (B) Time-course western blot analysis of pT72-RAB8A dynamics in BV2 cells following LLOMe treatment and washout. Cell lysates were probed with antibodies against pT72-RAB8A (top), total RAB8A (middle), and β-actin (loading control, bottom). (C) Representative immunofluorescence of acetylated α-tubulin (green), pT72-RAB8A (red), DAPI (blue) in WT and knockout BV2 cells after LLOMe (1mM, 30 min). Analysis confirms absence of primary cilia structures in these conditions. Scale bars: 5 μm (overview), 2 μm (insets). Data are representative of three independent experiments.

**Synopsis:**
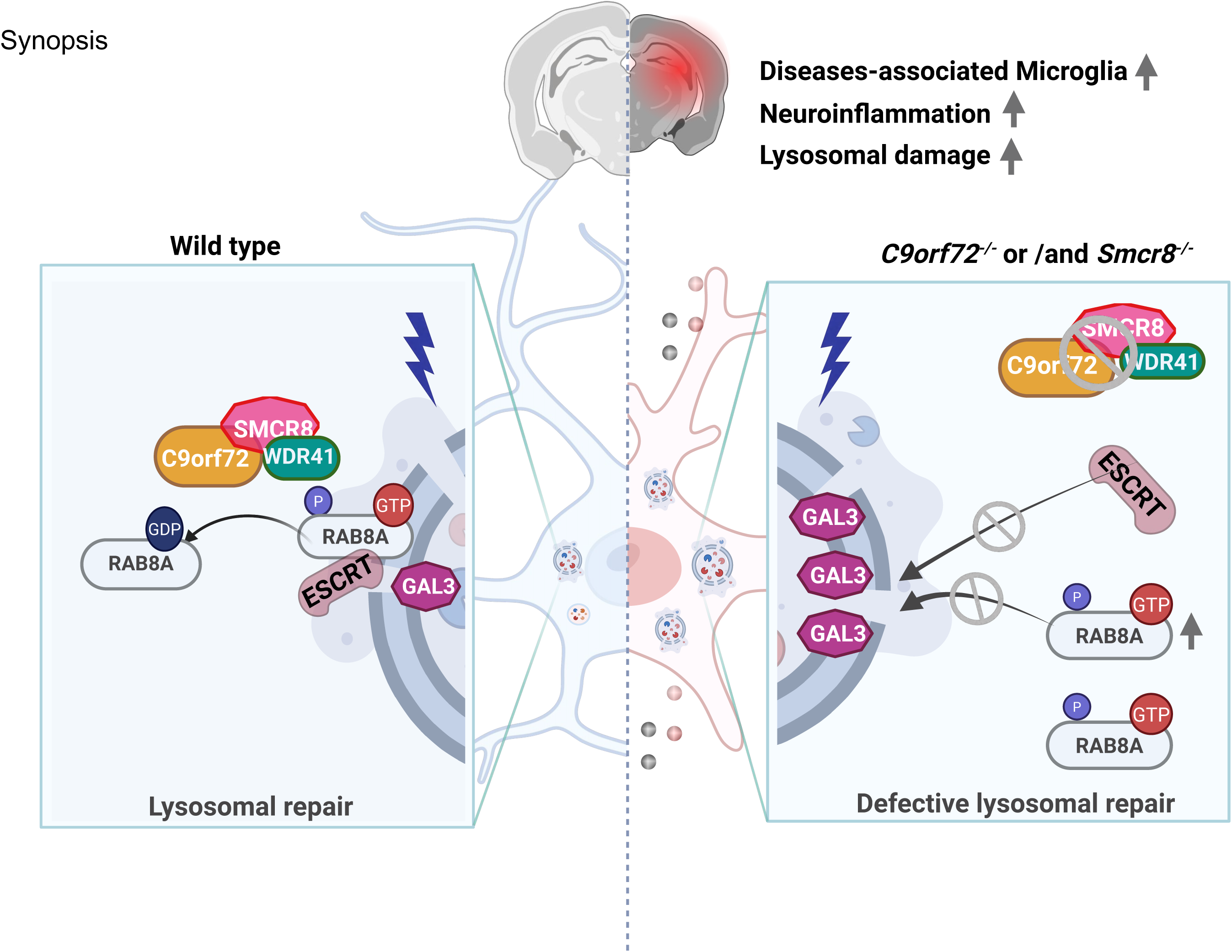
C9orf72/SMCR8 complex regulates lysosomal repair through RAB8A-ESCRT axis in microglia. Lysosomal integrity is safeguarded by the C9orf72/SMCR8 complex through regulation of RAB8A and ESCRT machinery. Upon lysosomal damage, phosphorylated RAB8A-GTP transiently localizes to lysosomes to recruit ESCRT for repair, and is subsequently inactivated by the GAP activity of the C9orf72/SMCR8 complex. Loss of the complex leads to accumulation of hyperphosphorylated RAB8A-GTP, defective ESCRT recruitment, persistent lysosomal damage, and induction of inflammatory disease-associated microglial states. WDR41 is included as a complex-stabilizing factor based on published interactions.

- Complex deficiency leads to lysosomal damage, altered DAM homeostasis, and increased microglial inflammation.
- Damaged lysosomes *in vitro* and *in vivo* are marked by increased Gal3 and p-RAB8A, serving as potential indicators of lysosomal injury.
- The C9orf72–SMCR8 complex regulates RAB8A GTP loading, which determines its phosphorylation, lysosomal targeting, and ESCRT-mediated repair.

